# Two-million-year-old microbial communities from the Kap København Formation in North Greenland

**DOI:** 10.1101/2023.06.10.544454

**Authors:** Antonio Fernandez-Guerra, Lars Wörmer, Guillaume Borrel, Tom O Delmont, Bo Elberling, Marcus Elvert, A. Murat Eren, Simonetta Gribaldo, Rasmus Amund Henriksen, Kai-Uwe Hinrichs, Annika Jochheim, Thorfinn S. Korneliussen, Mart Krupovic, Nicolaj K. Larsen, Rafael Perez-Laso, Mikkel Winther Pedersen, Vivi K. Pedersen, Anthony H. Ruter, Karina K. Sand, Martin Sikora, Martin Steinegger, Iva Veseli, Yucheng Wang, Lei Zhao, Marina Žure, Kurt H. Kjær, Eske Willerslev

## Abstract

Environmental DNA (eDNA) from the 2-million-year-old Kap København Formation of northern Greenland has revealed an ecosystem of plants and animals with no contemporary analogue^1^. Here, we reconstruct the microbial (bacterial, archaeal, and viral) communities that thrived at the site during this time. By leveraging a novel analytical framework that integrates taxonomic profiling, DNA damage estimates, and functional reconstructions, we identify and distinguish pioneer microbial communities from later permafrost microbial assemblages. We show that at the time of sediment deposition, the terrestrial input at the Kap København site originated from a palustrine wetland, suggesting warmer, non-permafrost conditions. During this period, the detection of methanogenic archaea and signals of their carbon metabolism is consistent with Kap København and similar northern ecosystems contributing moderate methane emissions. Intriguingly, we discover a remarkable nucleotide sequence similarity—exceeding 98%—between pioneer methanogens and present-day analogues in thawing permafrost. This aligns with the concept of “time-traveling” microbes^2^ surviving across geological time and waiting for conditions to turn favourable rather than evolving to adapt to changing conditions. Importantly, in contrast to the plant and animal communities of the Kap København, a striking similarity in microbial composition to that of a contemporary thawing Arctic suggests that microbial communities may serve as the first indication of broader climate-driven ecosystem disruptions.

## Main

Understanding how global warming will impact Earth’s biodiversity remains a subject of great debate. Some argue for tipping points where nature responds in drastic ways, reshuffling species distributions^3^, while others suggest a more gradual mode of change^4^. The predictability of biodiversity across time and space, i.e., whether species distributions, their compositions and ecosystem functions may repeat themselves across similar geological climate events, also remains contentious^5^. Addressing these issues is crucial for predicting future ecosystem transformations in response to climate change.

Of Earth’s many geological Epochs, the Pliocene to early Pleistocene (5 to 2 million years ago) is argued to be among the best analogues of a future world under global warming^6,7^. However, because of its time depth, the current understanding of biodiversity from this period is limited primarily to pollen^8^ and macrofossils^9^. Recent advantages in eDNA have pushed back ancient DNA retrieval from about 1 to 2 million years at the Kap København Formation of northern Greenland, a time when the average temperature was 10°C warmer than today^10^. The diverse plant and animal eDNA uncovered shows an environment with no modern equivalent, composed of taxa whose contemporary relatives are found in both Arctic and temperate environments^1^., implying a high degree of unpredictability in future species distributions under climate change.

Importantly, the eDNA study on the Kap København Formation did not consider the microbial communities. Microbes play a fundamental role in maintaining biogeochemical processes that influence microbial and non-microbial life^11–13^ composition and dynamics. Thus, characterising the ancient microbial eDNA component can provide a comprehensive view of the complex functioning of past ecosystems. Nevertheless, most ancient eDNA studies neglect the microbial communities because sustained microbial activity in sediments often disturbs and obscures the original microbial signatures^13^. Over time, the genetic signals of pioneering microbes—those present at the time of sediment deposition—become mixed with DNA from microbes that continue to thrive in the sediment, making it challenging to separate ancient microbial DNA from later microbial activity when reconstructing past ecological conditions and their impact on broader biogeochemical processes. As a result, ancient sediments contain two putative ancient DNA fractions^14^. One originates from organisms that died long ago and is highly fragmented and contains post-mortem modifications such as deamination damage at its termini ^15^. The other is DNA from organisms that have either remained dormant or survived with reduced metabolic activity for decades to hundreds of years (seed bank microbes) or for thousands to millions of years (“time-travelling” microbes). While seed bank microbes are considered part of modern ecosystems, acting as a genetic reservoir, “time-travelling” microbes can reintroduce ancient genetic traits into present-day ecosystems^16^.

For this reason, DNA from seed bank and “time-travelling” microbes is challenging to distinguish from modern contaminant DNA. To be able to analyse ancient microbial diversity from eDNA records, we developed a new analytical framework that integrates taxonomic profiling, DNA damage estimates, and functional reconstructions and applied it to the Kap København Formation.

### The Kap København Formation

The Kap København Formation consists of a c. 100 m thick succession of shallow marine sediments primarily accumulated during a ∼20,000 years long interglacial period about two million years ago^1,17^. These sediments serve as an archive for capturing the community composition upstream from the depositional sites, as they were fluvially transported to the foreshore and concentrated as organic detritus mixed into sandy near-shore sediments. After the Kap København Formation was deposited, it was uplifted due to the solid Earth’s flexural isostatic response to erosional unloading by fjord incision^18^ (Fig. 1B). We estimate an uplift rate of ∼0.09 mm/yr, which raised these shallow marine deposits to their current elevation (Fig. 1C; Supplementary Table S3). Based on this rate, we infer that the four sampled localities (localities 50, 75, 119, 69) (Fig. 1B) across three stratigraphic units (B1, B2, and B3; from older to younger) were exposed subaerially between 0.8-1.2 million years ago (Fig. 1C). Sediment records from below the present ice sheet suggest that the Greenland Ice Sheet was already established at the time of subaerial exposure 1.2 million years ago^19^. It is reasonable to assume that during the period when the sediments emerged subaerially, they gradually became part of the permafrost, creating a distinction between the pioneer communities and the microbial communities that have inhabited the permafrost sediments up to the present day (Fig. 1C).

**Fig. 1.**
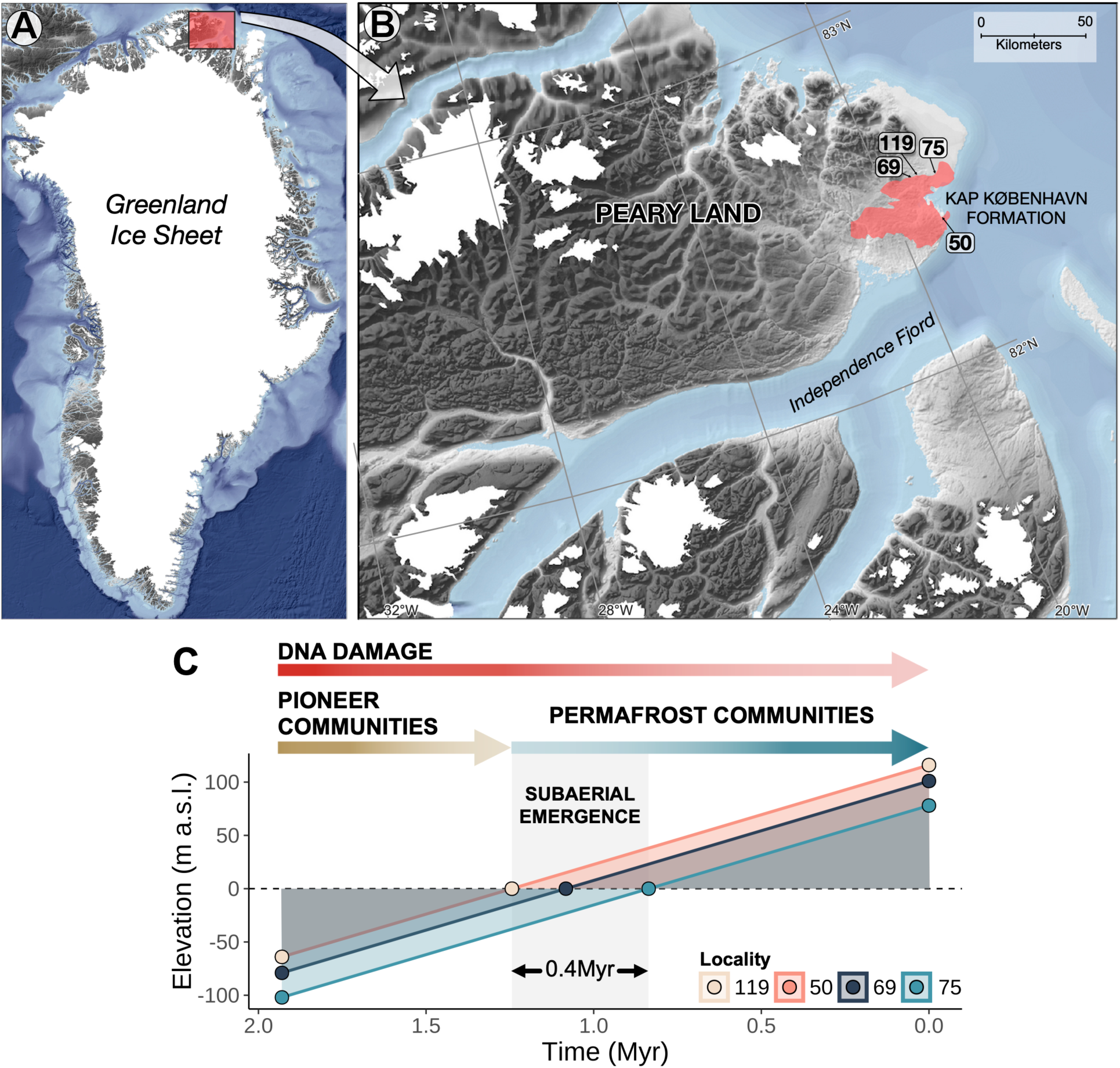
Geographical location and uplift model. A, Map indicating the location of Kap København Formation in North Greenland at the entrance to the Independence Fjord (82° 24′ N 22° 12′ W). B. Spatial distribution of the ∼100-m thick succession of shallow marine near-shore sediments between Mudderbugt and the low mountains towards the north, circles highlight the localities used in the study. C, Simple uplift model of the Kap København Formation. The darker colour in the DNA damage arrow indicates higher damage. In contrast, the arrows for the pioneer and permafrost microbial communities show abundance levels, with darker colours indicating higher abundance. Localities 119 and 50 are superimposed.

### Pioneer and permafrost microbial communities

Most endogenous DNA in ancient sediments is highly fragmented and subject to deamination damage at the termini. Therefore, we designed a novel approach for inferring microbial community composition in deep-time samples from ultra-short and damaged DNA sequencing reads. Our approach involved three main components. Firstly, we created a comprehensive genomic database with a common taxonomic framework (Extended Data Fig. 1) covering Archaea, Bacteria, Viruses and eukaryotic organelles. Secondly, in our workflow (Extended Data Fig. 2), we performed a sensitive search using Bowtie2^20^ and applied stringent filtering criteria that leveraged read patterns across the references, allowing us to confidently retrieve low-coverage genomes, even with as little as 1% of their genome present (hereafter, we will refer to the “breadth of coverage” as detection, the proportion of a given genome that is covered at least 1X). We inferred their relative abundance by normalising the estimated TAD80^21^ (truncated average depth) number of reads to the reference length. Lastly, we performed large-scale damage estimation on the filtered results to identify those references that are potentially members of the pioneer microbial communities. We excluded taxa identified in the controls and blanks (Supplementary Table S2; Extended Data Fig. 3), as well as the eukaryotic hits from subsequent analyses, since the eukaryotic references were intended to be used as bait in the competitive mapping and as a reference to select the damage threshold (Extended Data Fig. 4A). We then focused our analyses on 22 samples across the stratigraphic units (B1, B2, and B3) (Supplementary Table S1) that met two criteria: (1) having more than 10 million unique reads and (2) where the combined relative abundance of references exhibiting damage greater than 0.11 (we used the background-subtracted damage frequency at the first position of the reads provided by metaDMG^22^, where one represents the maximum level) accounted for at least 50% (Extended Data Fig. 4B). We mapped an average of 7.2% of unique reads per sample with an average of 1,950 references (Fig. 2A; Supplementary Table S4). Finally, we aggregated our results by calculating the geometric mean of taxonomic abundances and averaging the inferred damage at the species level. On average, the damaged fraction composed 84% of the estimated relative abundances in each sample, revealing Archaea as the dominant pioneer group (Fig. 2A; Supplementary Table S4).

**Fig. 2.**
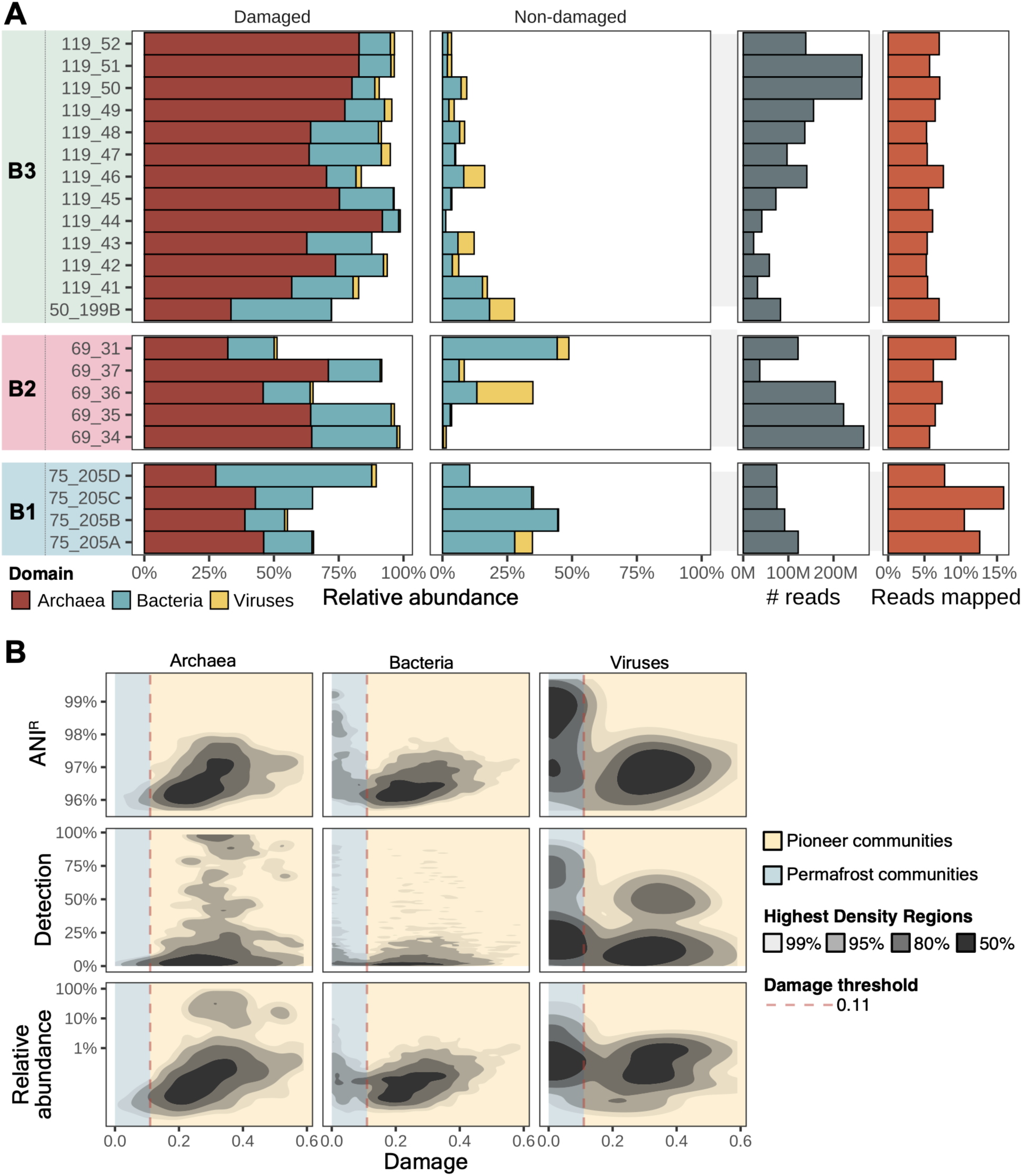
Damage and non-damage distribution patterns. A, Proportion of taxa identified as damaged and non-damaged in the three stratigraphic units B1 to B3 of the Kap København Formation based on the estimated taxonomic abundances. The two rightmost panels show the number of dereplicated reads used for mapping and the percentage of reads successfully mapped to any references within the database employed for taxonomic profiling. B, The Highest Density Regions plots show the average nucleotide identity, detection, and relative abundance distribution as a function of the damage for Bacteria, Archaea and Viruses. The dashed red line sets the damage threshold we inferred from the Eukaryotic data. The potential fraction of pioneer communities is highlighted in yellow, while the potential contemporary communities inhabiting the permafrost are marked in blue.

By integrating damage estimates, relative abundances, detection, and mean read Average Nucleotide Identity estimates (ANI^R^), we differentiate between the initial pioneer communities and those that inhabited the permafrost after the subaerial emergence. Our results indicate a limited presence of Archaea in the permafrost communities (blue area in Fig. 2B), consistent with the low abundance of archaeal taxa typically found in the cryosphere^23,24^. On the contrary, the high values for ANI^R^, detection and relative abundance of the bacterial and viral fraction indicate the distinct composition of contemporary permafrost communities. Despite the significant time gap between the modern reference genomes and the sequences identified in the pioneer communities, we observed elevated ANI^R^ values even when considering the extent of damage in the sequences. In some cases, the ANI^R^ values approached 98%, indicating a relatively high level of identity. Remarkably, our samples show high detection values for Archaea and some bacterial taxa, where some of them have more than 75% of the genome recovered (Fig. 2B; Supplementary Table S4).

### Modern analogues to the Kap København Formation

After disentangling the pioneer from the permafrost microbial communities, we investigated potential modern analogues of the pioneer microbial communities. Firstly, we employed meta-Sourcetracker^25^, trained on over 1,003 modern metagenomes, to identify the contribution of 33 different biomes to our samples (Supplementary Table S8). We further supplemented our analysis by employing a k-mer similarity method^26^ on the reads retrieved from taxa identified as part of the pioneer communities. In both cases, the *Aquatic:Freshwater:Sediment* biome was the most significant contributor to all samples, with proportions as high as 80% in some cases (Fig. 3A and Supplementary Table S9). The second most important contributor was *Terrestrial:Source:Permafrost*, with proportions higher than 25% in some samples. The biome *Terrestrial:Soil:Forest soil* was the third most significant contributor, with some samples having a proportion of 10%. While meta-Sourcetracker failed to detect the marine signal in some of our locations, the k-mer-based method showed higher sensitivity, identifying a 10% contribution from the biome *Environmental:Aquatic:Estuary* in a sample from location 75 (Supplementary Table S9).

**Fig. 3.**
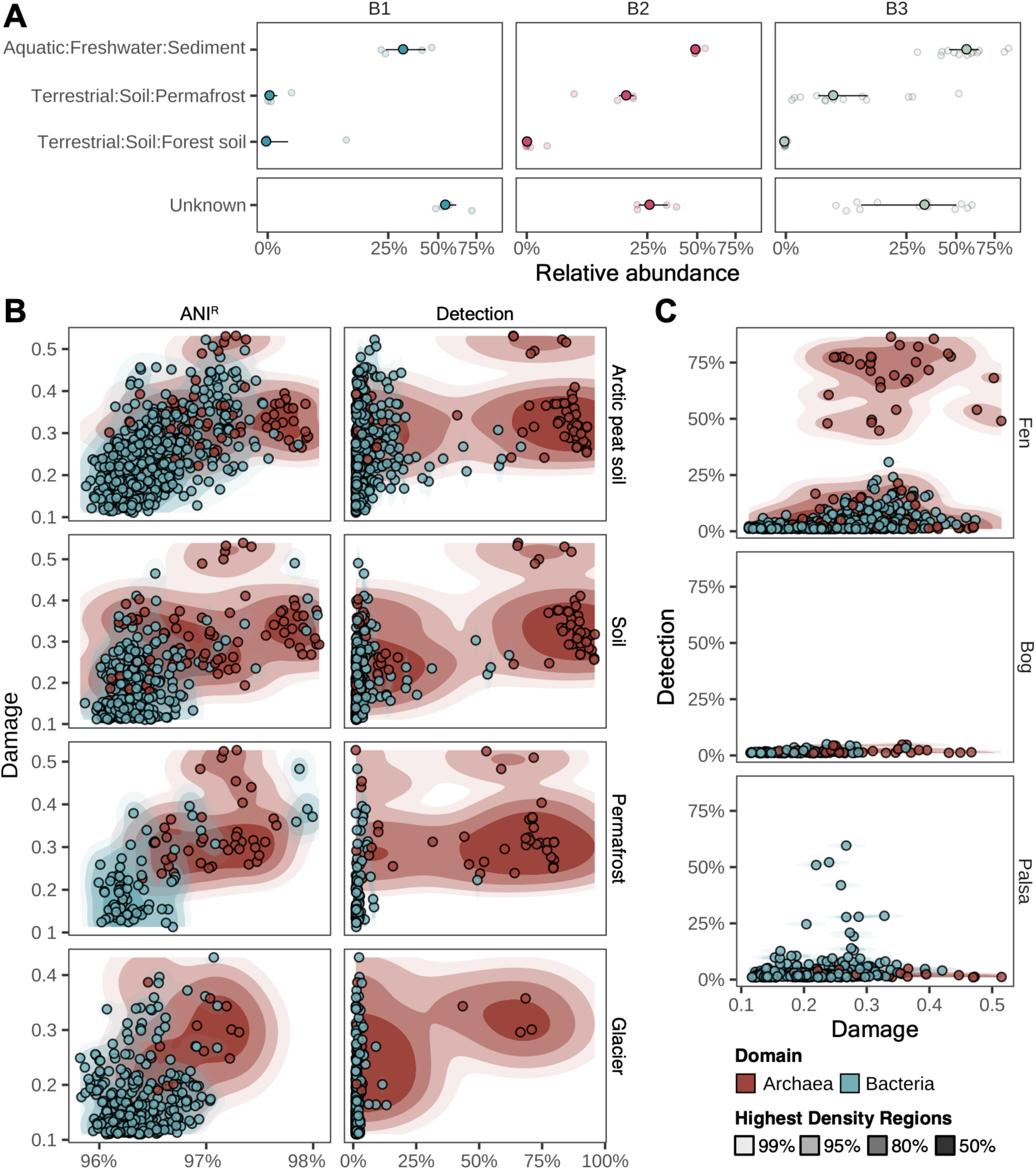
Kap København Formation modern analogue biomes. A, Boxplot representing the estimated proportions by meta-SourceTracker from each “source” or biome for a single Kap København Formation sample or “sink”. Colours correspond to the three stratigraphic units, B1, B2 and B3. B, Highest Density Regions plots depicting the relationship between damage and the average nucleotide identity, as well as the detection (the proportion of the reference that is covered) after mapping reads from the damaged taxa against metagenome-assembled genomes recovered from the biomes described in Nayfach et al. 2020^27^ and Liu et al. 2022^23^. C, Detection and damage of the recruited genomes through mapping the reads from damaged taxa against metagenome-assembled genomes obtained from a thawing permafrost gradient.

Next, we explored the distribution of the reads extracted from the pioneer communities in each sample using the Genomes from Earth’s Microbiomes (GEM) catalogue^27^, which covers diverse habitats of Earth’s continents and oceans and the glacier microbiome catalogue^23^. We used the estimated damage, ANI^R,^ and detection to select the biomes that recruited multiple non-unique references with a detection rate higher than 50%. The biomes with the highest number of references (Fig. 3B) were consistent with the results from the microbial source tracking methods, with archaea having a detection rate of over 75% and a high ANI^R^ despite high levels of estimated damage (Supplementary Table S10). The Arctic peat soil biomes and soil had the highest number of references, indicating the successful recovery of the pioneer microbial communities. To increase the resolution of our analysis, we calculated where our samples would fall on a permafrost thaw gradient (Supplementary Table S11). We used the metagenome-assembled genomes recovered from the Stordalen Mire^28^ to recruit the reads from the pioneer communities. Our results show that fens, which are peat-forming wetlands, recruit the largest number of archaeal MAGs with detection above 50% (Fig. 3C). We also found microbial genomes with moderate detection in palsa (Fig. 3C), which aligns with the permafrost signal detected by both the microbial source tracking methods and the read recruitment to genome collections.

It is worth noting that the samples collected from the Kap København Formation aggregate diverse microbial communities sourced from upstream locations mixed with *in-situ* micro-environments within the delta, as observed in extant delta systems^29^. These communities have been deposited over 20,000 years, resulting in a blend of local and upstream contributions reflecting the depositional history of the formation. This composite estimate serves as a representation of the dominant biome during the Early Pleistocene period. Furthermore, the vegetation recovered on the site reinforces our findings that the region experienced prolonged periods with annual mean temperatures well above freezing and consequently without permafrost (Supplementary Information, section 1).

### Community structure of a two-million-year-old microbiome

The structure and composition of microbial communities preserved in sediments from two million years ago revealed remarkable similarities to those in modern palustrine wetlands. Reconstructing the abundance of such ancient microbiomes presents significant challenges due to the taphonomic and biological processes that DNA undergoes over geological timescales^30,31^. Nevertheless, we identified a diverse array of archaeal and bacterial taxa from our dataset (Supplementary Table S4). Our analyses suggest that methanogenic archaea were the dominant group at the Kap København site two million years ago (Fig. 4A). Considering the warmer conditions of the Early Pleistocene, the presence of methane-producing microbial communities upstream of the deposition site two million years ago resembling those found in modern wetlands or thawing permafrost, is conceivable. However, their unexpectedly high relative abundance (averaging over 60% in most samples) is unlikely. While methanogens can show higher abundances than other archaea in a thawing permafrost environment^32^, they typically account for only 1-5%^33–37^ of the overall community in present-day settings. Methanogens rely on the by-products of bacterial metabolism for energy, but bacteria consume most of the energy available in organic matter, leaving only a small fraction for methanogenesis^38^. To discard potential DNA extraction biases, we validated our results using a new optimised extraction protocol in a subset of our samples (Supplementary Information, section 2; Supplementary Table S5-S6; Extended Data Fig. 5). Multiple factors might contribute to the increased preservation of archaea. Firstly, differences in the local chemical environment, influenced by the metabolic activity of archaea, could contribute to preservation disparities. Secondly, most archaea possess distinct cell envelopes consisting of a lipid membrane externally covered by a paracrystalline surface (S-) layer composed of one or two proteins with strong inter-subunit interactions^39^. Several features of the S-layers could be responsible for the increased preservation; the S-layers can repair themselves against damage and are also found to serve as nucleation sites for mineral formation, resulting in encrustation^40^. Finally, differences in chromatin structure could also influence preservation. DNA is complexed with histones in many archaea, including methanogens, making them among the most abundant proteins in *Methanobacteriales*^41^. This distinct chromatin organisation in archaea compared to bacteria may be yet another factor contributing to the observed differences in preservation patterns.

**Fig. 4.**
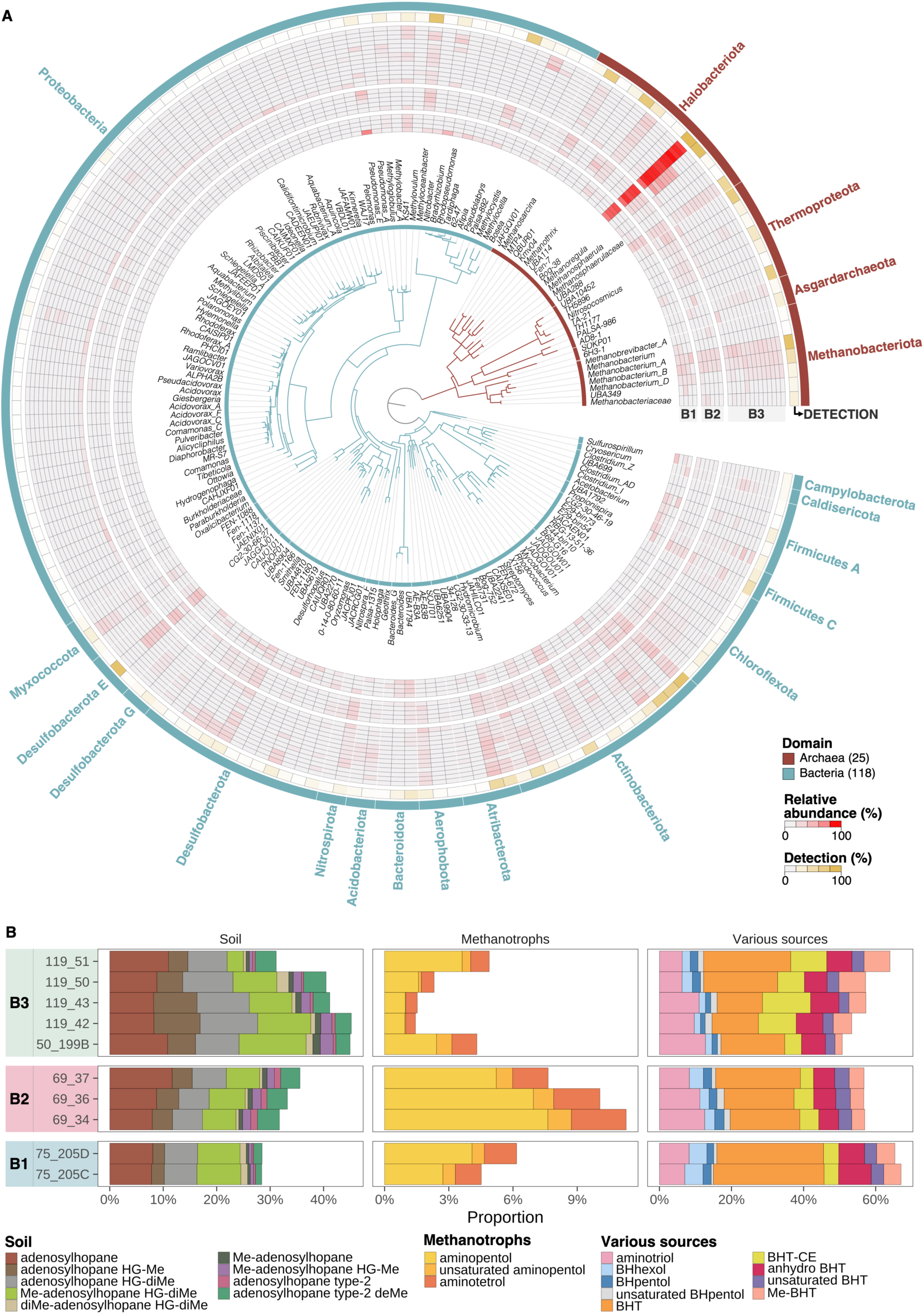
Community composition of the archaeal and bacterial taxa. A) Taxonomic profile of the damaged identified taxa, mapped into the GTDB phylogenomic tree at the genus level. Only genera that make up at least 1% of the taxonomic abundance at the family level are displayed. Each layer of the circle corresponds to a different sample grouped by member unit. The coloured sections of the circle indicate the relative abundance of each genus in the sample, with the intensity of the colour representing the transformed square root of the proportion. The outer layer corresponds to the maximum detection value for the references in a genus across samples. B) The relative abundance of bacteriohopanepolyol derivatives is expressed in percentages of the total. Adenosylhopanes are named after Hopmans et al.^42^.

We identified a variety of methane-related archaea across multiple genera, each showing detection rates of 75% or higher. These include the CO_2_-reducing hydrogenotrophic g Bog-38 (*Methanoflorens*), *Methanoregula*, and *Methanobacterium_A*. We also detected the acetoclastic methanogen *Methanothrix*, as well as the versatile methanogen *Methanosarcina* (Fig. 4A). Additionally, we observed the archaeal methanotroph g Kmv04 (*Methanocomedenaceae*). Our analyses also revealed the presence of bacterial phyla such as *Atribacterota* (g CG2-30-33-13) and *Aerophobota* (g AE-B3A), which are commonly found in methane-rich environments ranging from temperate soils and deep marine sediments to permafrost^43,44^. Moreover, our study identified a range of methanotrophic bacteria exhibiting high detection rates. Among these, we found high-affinity methanotrophs such as *Methyloceanibacter*^45–48^, as well as genera within the *Methylomonadaceae,* such as *g KS41*, commonly found in acidic forest soils^49^, and *Methyloglobulus*, usually found in lake sediments^50^.

We further supported the patterns recovered from ancient DNA by analysing microbial lipid biomarkers and their carbon isotopic compositions for a subset of our samples. This data provided additional compelling evidence for a dynamic microbial turnover of methane by detecting lipids from methanogens and methanotrophs. Among the archaeal lipids, we identified a predominance of archaeol and hydroxy-archaeol^51^ (Supplementary Table S7), which are specifically abundant in methanogenic archaea, particularly *Methanothrix* and *Methanosarcina*^51^. Carbon isotopic analysis for archaeol in sample 69_34 revealed a δ^13^C value of -21.6‰, consistent with CO_2_ and/or acetate as a carbon source for lipid biosynthesis (Supplementary Information, section 6). The detection of GDGT-0 further corroborates the presence of hydrogenotrophic methanogens, aligning with the DNA-based results.

Additionally, the lipid pool showed a notable abundance of bacteriohopanepolyols (BHPs) derived from soil bacteria, increasing from 28% in B1 to 41% in B3 (Fig. 4B). This is an indicator of a marked input of soil organic matter, consistent with fluvial deposits originating from permafrost environments as well as forest soils^52^. In unit B2 (Fig. 4B), the high relative proportion of typical methanotrophic BHPs, such as aminotetrol and aminopentol, combined with a distinctly ^13^C-depleted signal observed in the hopanoid diplopterol (hopan-22-ol) with a δ^13^C value of -61.6‰, points towards localised hotspots of methane oxidation by aerobic methanotrophic bacteria upstream. Such methanotrophic activities potentially contributed to reducing methane emissions in high-latitude ecosystems (Supplementary Information, section 3) despite the northward expansion of wetlands.

We also identified microorganisms involved in carbon and nitrogen cycling processes. These include members of the families *Bacteroidaceae* (g UBA1794 and *Bacteroides*) and *Clostridiaceae* (*Clostridium_AD*); the phylum *Myxococcota* (g Fen-1088, found in symbiotic relationships with the arbuscular mycorrhizal fungal hyphae^53^), and members of the genus *Propionospira*, which produce propionate, acetate, and CO_2_ as the main products of their carbohydrate fermentation^54,55^. We also detected predicted homoacetogenic members of the family f UBA5619 (g UBA5619), which may contribute fermentation products to *Methanothrix* while removing H₂ and formate from the environment^56^. *Eubacteriaceae* (g UBA1792), capable of degrading substrates such as fructose and mannose (Supplementary Table S4), and g JADGOW01, possessing the necessary gene repertoire for acetogenesis (Supplementary Table S4), were also identified. We additionally observed members from the family *MBNT15* (g CG2-30-66-27), commonly found in peatlands, where they act as scavengers by fully mineralising small organic molecules produced during the microbial breakdown of complex polymeric compounds^57,58^. We detected ammonia-oxidising archaea (AOA) from the *Nitrososphaeraceae* (g UBA10452 and g TA-21), which are found in acidic polar and alpine soils^59^ and a High-Arctic glacier^60^, respectively, and may indicate the presence of aerobic niches. Besides the AOA, we also identified bacterial taxa involved in nitrogen fixation, including *Pseudolabrys* (73% detection rate) and *Bradyrhizobium*, as well as comammox bacteria from *Nitrospiraceae* (g Palsa-1315) capable of complete ammonia oxidation^61^. Finally, we identified the archaeal *Thermoproteota* genus g AD8-1 (70% detection rate) from the class *Bathyarchaeia* and the *Asgardarchaeota* genus g 6H3-1 from the class *Lokiarchaeia*, both of which inhabit marine sediments. The closest reference from *g AD8-1* to our data was reconstructed from permafrost samples with marine influence^62^. In contrast, the closest reference from g 6H3-1 was isolated from deep-sea sediment samples of the Hikurangi Subduction Margin^63^.

### Microbial carbon processing

Our taxonomic profiling analysis (Fig. 4A) shows that the dominant group consists of CO_2_-reducing hydrogenotrophic members and other versatile methanogenic archaea. Furthermore, our results revealed the presence of high-affinity methanotrophic bacteria in the samples. We complemented these taxonomic analyses with ecosystem-wide metabolic reconstructions^64^ to identify the primary metabolisms associated with methanogenesis two million years ago.

Peatlands store vast amounts of carbon from plant polymers such as cellulose and hemicellulose. In present-day peatlands, these polymers are degraded by microorganisms, forming glucose, xylose, and N-acetylglucosamine. These compounds are further converted into low-molecular-weight alcohols and organic acids such as ethanol, propionate, acetate, and lactate, as well as H_2_ and CO_2_ via fermentation. These fermentation products serve as substrates for CO_2_-reducing hydrogenotrophic and acetoclastic methanogens, which utilise H_2_/CO_2_ and acetate, respectively^65^. Given the ubiquity of these processes in modern peatlands, we expected to observe evidence of similar metabolic pathways in our ancient samples. We compiled a collection of standard and custom KEGG modules and specific CAZymes^28^ (Carbohydrate-active enzymes) to reveal the different steps involved in carbon processing by the pioneer communities. Our analysis showed evidence of all carbon processing steps in our samples (Fig. 5, Supplementary Table S12-13), even when only using reads that map to the pioneer communities (Fig. 5, tiles with thicker borders; Supplementary Table S14-15). We applied a highly conservative approach by using translated searches with ultra-short damaged reads and only considering those modules with completion over 80%. In agreement with the taxonomic profiling, we identified pathways for CO_2_-reducing hydrogenotrophic, acetoclastic, and methylotrophic methanogenesis, as well as aerobic methanotrophy (Fig. 5). The co-occurrence of this pathway suggests that a significant proportion of CH4 might have been oxidised, thereby limiting emissions to the atmosphere two million years ago^47,66^. Our results may underestimate the true functional potential of the ancient microbial community due to cytosine deaminations, which can introduce stop codons and non-synonymous substitutions in the predicted open reading frames used in our translated searches. Our stringent pathway identification thresholds, coupled with the presence of deaminations, likely contributed to the failure to identify methane pathways in B1. Indeed, the average damage values for the methanogenic archaea in B1 were 0.48 ± 0.05, amongst the highest values estimated in our dataset, a pattern also observed in the eukaryotic fraction. However, we detected methane pathways in some sites from B1 (acetoclastic methanogenesis in 75_205B and CO_2_-reducing hydrogenotrophic methanogenesis in 75_205A) by lowering the pathway identification threshold to 0.7.

**Fig. 5.**
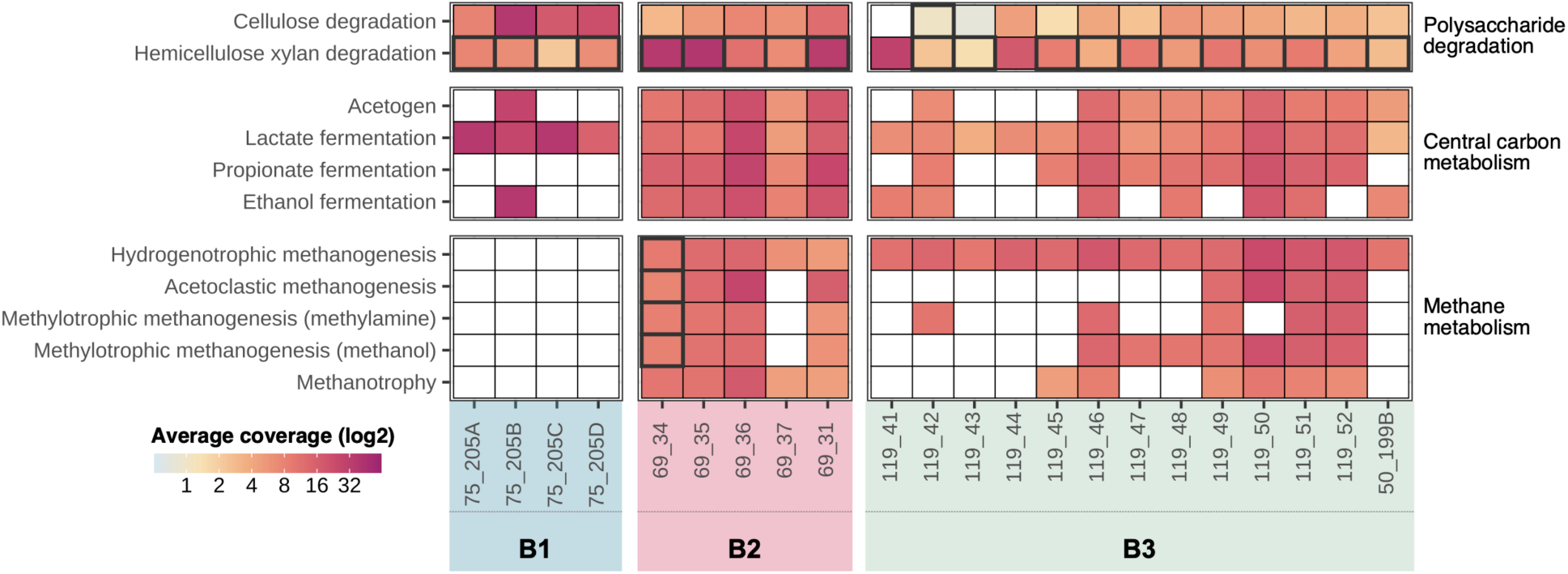
Carbon metabolism in the Kap København Formation. Functional profile depicting pathways involved in the carbon metabolism in the permafrost. Tiles with thicker borders represent samples where the pathways were also detected using only reads that map to damaged references. White tiles indicate samples where no pathways have been detected.

### The Kap København Formation virome

Viruses play a major role in structuring microbial communities across ecosystems^67^; hence, we further explored the community structure of the Kap København Formation by characterising its DNA virome. We recovered the reads mapping to references from the IMG/VR included in our database from the taxonomic profiling. To ensure the accuracy and reliability of the results, we only considered the references with a detection rate above 10%. We recovered 36 references, 15 damaged and 21 non-damaged (Fig. 6A), all belonging to the class *Caudoviricetes* (phylum *Uroviricota*), the dominant group of prokaryotic viruses (Supplementary Tables S4, S16). To further expand the reach of the homology detection, we searched the translated reads against the proteomes of viruses from the IMG/VR included in our database and a curated collection of mobile genetic elements (MGE) associated with methanogenic archaea^68^. After the search step, we identified which proteins were more likely to be present in a sample and then selected those genomes with over 20% of their proteins detected. This approach detected 231 reference viruses, establishing a detailed view of the two-million-year-old virome (Supplementary Tables S4, S16). Although members of the *Caudoviricetes* again represented the dominant (70%) component of the virome (Fig. 6B), a considerable fraction could be affiliated with the realms *Monodnaviria* (viruses with ssDNA genomes) and *Varidnaviria* (non-tailed icosahedral viruses with double jelly-roll capsid proteins). Notably, the identified monodnaviruses belong to three different phyla specific to eukaryotic (*Cressdnaviricota*) and bacterial (*Hofneiviricota* [filamentous phages] and *Phixviricota* [phiX174-like phages]) hosts. Members of the *Cressdnaviricota* represent some of the simplest viruses associated with diverse eukaryotic hosts, including protists, fungi, plants and animals. Additionally, *Varidnaviria* was represented by viruses of the class *Megaviricetes* (phylum *Nucleocytoviricota*), which includes environmentally widespread eukaryotic viruses with big and giant dsDNA genomes. Finally, the identified methanogenic MGE, which include members of the *Caudoviricetes*, archaea-specific lemon-shaped viruses and several unclassified MGE (Fig. 6C), are associated with *Methanosarcinales*, *Methanobacteriales* and *Methanomicrobiales*, all of which were detected in our samples (Fig. 4A).

**Fig. 6.**
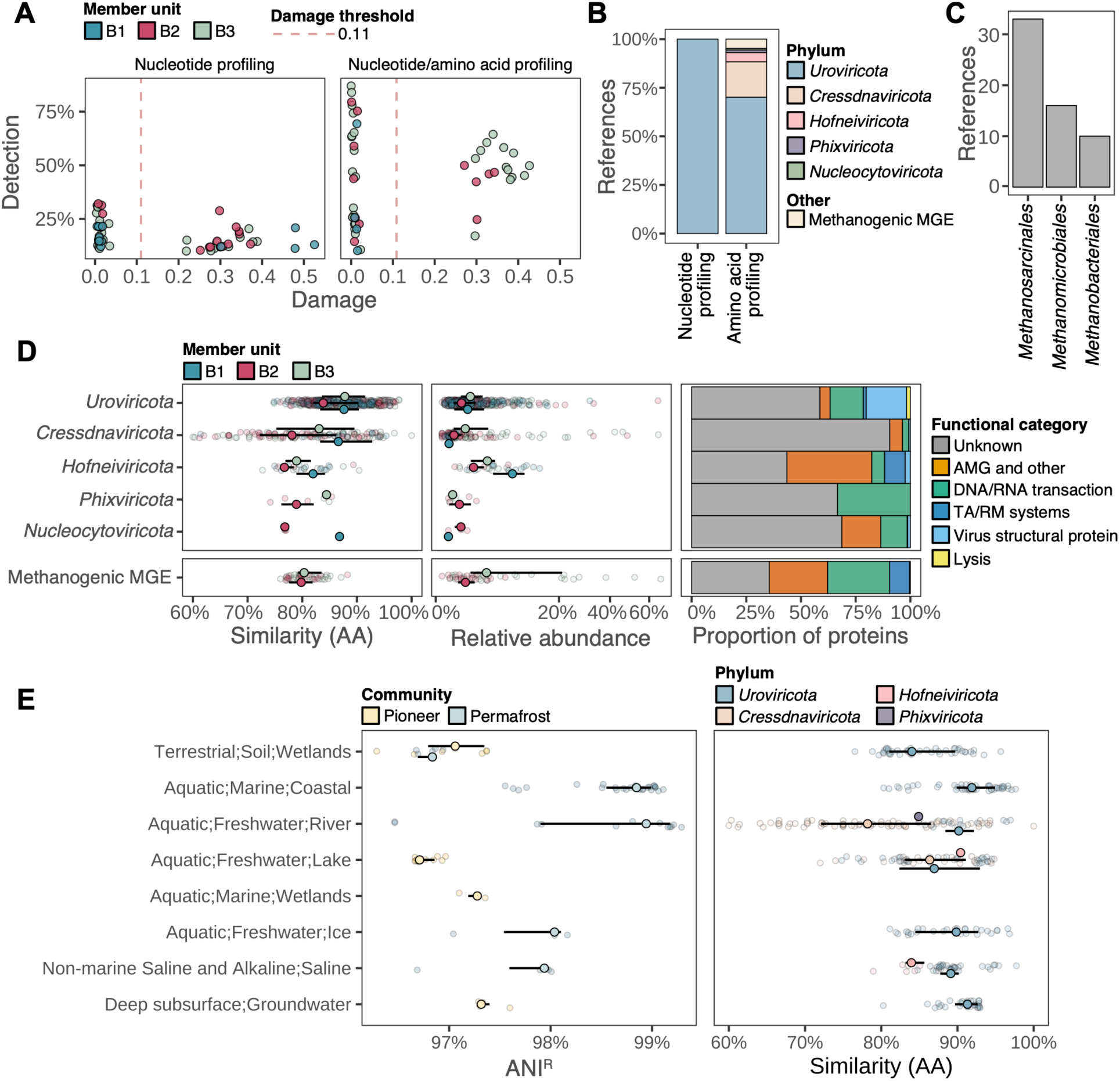
The Kap København Formation virome. A, Each point depicts a viral reference identified in a sample by nucleotide profiling on the left panel and by nucleotide and amino acid profiling on the right panel. Red dashed line depicts the damage threshold we used to delimit the pioneer from the permafrost communities. B, Proportion of references detected in the whole dataset by phylum and methanogenic mobile genetic elements. C, Number of references associated with each predicted host for the methanogenic MGEs. D, The first two panels show the distribution of amino acid similarity values and inferred abundances for each viral reference identified in each sample. The third panel corresponds to the function categories of the proteins annotated to each group, phylum or methanogenic MGE. E, Environmental distribution of the different references we recruited provided by the ecosystem information in the IMG/VR v4 database

On average, the viral fragments we retrieved exhibited an amino acid similarity (PAM30) of 84% to the references (Fig. 6D). Among them, the members of *Cressdnaviricota* displayed the largest divergence from modern references, with similarities as low as 60%. These similarity values indicate a significant divergence of the viruses found in the formation compared to modern references. These viruses would have gone undetected if only nucleotide assignment methods were employed. Both approaches identified 14 *Caudoviricetes* references (Fig. 6A), with three of them showing substantial detection (>50%) and damage (>0.3). Considering the amount of damage, these virus genomes show a high identity to the reference, with ANI^R^ exceeding 96%. Particularly noteworthy is the similarity to the reference IMGVR_UViG_2617271244_000001 (detection: 64%, damage: 0.34) with an ANI^R^ of 97% and an amino acid similarity of 95%. This virus is predicted to infect *Protochlamydia naegleriophila*^69^, an obligate intracellular symbiont of the free-living amoeba *Naegleria*, commonly found in warm freshwater^70^. *Caudoviricetes*, *Cressdnaviricota* and the methanogenic viruses have the highest relative abundance, with certain samples surpassing 20% of the total viral fraction (Fig. 6D, Supplementary Tables S4, S16).

The amino acid searches also provided information about the functional potential of the viruses we recruited (Supplementary Table S17). Although a large proportion (71%) of the recruited viral proteins lack functional annotation, the remaining 29% of proteins validated the taxonomic assignment of the corresponding reads and provided valuable insights into the functional capacity of the virus community (Fig. 6D, Supplementary Table S17). In particular, the two largest functional categories corresponded to taxon-specific virus structural proteins (16%) and proteins involved in DNA/RNA transactions (genome replication, transcription, recombination; 9.5%), respectively. We also identified proteins responsible for host cell lysis and toxin-antitoxin and restriction-modification systems. Finally, viruses encoded diverse auxiliary metabolic genes involved in various metabolic pathways (phosphoheptose isomerase, UDP-glucose dehydrogenase, phosphoadenosine phosphosulfate reductase, sulfatase-modifying factor enzyme, deoxyribonucleoside 5’ monophosphate phosphatase, nitroreductase, glutamine amidotransferase, cation efflux protein), enzymes for modification of various substrates (acetyltransferase, nucleotide kinase, serine-threonine kinase, GtrB glycosyltransferase for O-antigen conversion), hydrolases (glycoside hydrolase, esterase/lipase, proteases) as well as proteins responsible for modulation of host responses (MazF-like growth inhibitor, sporulation stage III protein D, translational regulator).

We leveraged the ecosystem information associated with each reference in the IMG/VR data to gain insights into the environmental origins of the references identified through both approaches (Fig. 6E). Consistent with the source tracking analyses, most of the references originate from aquatic freshwater biomes. Notably, the references with the highest ANI^R^ and amino acid similarity are derived from environments that resemble permafrost conditions. The biome labelled Aquatic;Marine;Coastal corresponds to samples obtained from arctic subzero sea-ice and cryopeg brines, while the Non-marine Saline and Alkaline;Saline biome represents samples collected from Lost Hammer Spring in Axel Heiberg Island situated in the Arctic Ocean, as well as various lakes in Antarctica.

### A “time-travelling microbe” at the Kap København Formation

The remarkable sequence similarity between the pioneer microbial communities in Kap København and modern references from permafrost thawing gradients^28^ offers an exceptional opportunity to gain ecological and evolutionary insights from our data. Members of the genus *Methanoflorens* (*g Bog-38*), which had the highest read recruitment level in our study at high identity (>98%) despite the presence of damage (median damage across samples of 0.3), represent a fascinating case. *Methanoflorens* is a CO_2_-reducing methanogen which is widely distributed in high methane-flux habitats. In the context of thawing permafrost, *Methanoflorens* shows a high abundance compared to other methanogenic microorganisms and is a significant contributor to methane production^32^. The *Methanoflorens* reference 3300025461_7, which recruited most of the reads in all samples of our study, is a metagenome-assembled genome derived from a drained thaw lake basin in the Arctic coastal plain near Barrow, Alaska^71^.

We decided to test whether the reads showing >98% similarity to contemporary *Methanoflorens* from thawing permafrost could originate from “time-travelling” microbes^16^. Each of our samples represents an isolated microcosm that has undergone multiple environmental transitions, beginning as terrestrial sediment accumulating in the delta, passing through shallow marine sediments, and ultimately becoming permafrost for the past million years following the uplift of the formation (Fig. 1C). If these reads indeed derive from “time-travelling microbes”, we expect to find a mixture of both damaged and undamaged DNA fragments from the same *Methanoflorens* lineage, spanning different periods. The pioneer communities, those from the deepest past, should have clear post-mortem damage signals, such as shorter fragment lengths and deamination patterns. In contrast, fragments from organisms that have remained dormant should lack these damage patterns and be longer.

To disentangle the signal of these pioneer methanogens (reads with post-mortem damage) and those who survived in the permafrost (reads without post-mortem damage), we analysed the data collected from sample 119_50, where the genome with the accession 3300025461_7 successfully recruited 1,114,032 reads (estimated damage: 0.26, detection: 94% and ANI^R^: 98.2%). We used a novel method^72^ to calculate the probability of the read being ancient based on the information on nucleotide changes and the fragment length. To ensure accurate inference of the probabilities of being ancient, we initially excluded from our analysis reads that exhibited a 100% identity to the reference and had a read length smaller than 40 nt (modal read length + 5nt) from our analysis.

This precautionary step minimised potential interferences caused by a skewed distribution of identical reads (Fig. 7A). An assumption of the method is the requirement for prior knowledge of the modern contamination rate in the sample. Given the inherent difficulty in estimating this rate for our sample type, we adopted a pragmatic approach. We employed modern contamination rates of 5% and 10% and then calculated the average of the resulting probabilities. The results of the inference highlight that we have a group of reads that have a high probability of being ancient (AN) and damaged (PD) and a second group that has a low probability of being damaged but a high probability of being ancient due to their read length (Fig. 7B, left panel). The pioneer fraction exhibited ***λ***=0.44, ***δ***d=0.029, ***δ***s=1 and ***v***=0.11. On the other hand, the surviving fraction in the permafrost had ***λ***=0.23, ***δ***d=0, ***δ***s=0, ***v***=0. The larger values of ***δ***d, ***δ***s, and ***v*** and smaller values of ***λ*** typically indicate a more severe level of DNA damage. Thus, Briggs parameters suggest that the pioneer fraction shows clear indications of damage, unlike the permafrost fraction, confirming the ancient nature of reads from the sample 119_50 that matched the reference 3300025461_7 with high levels of identity.

**Fig. 7.**
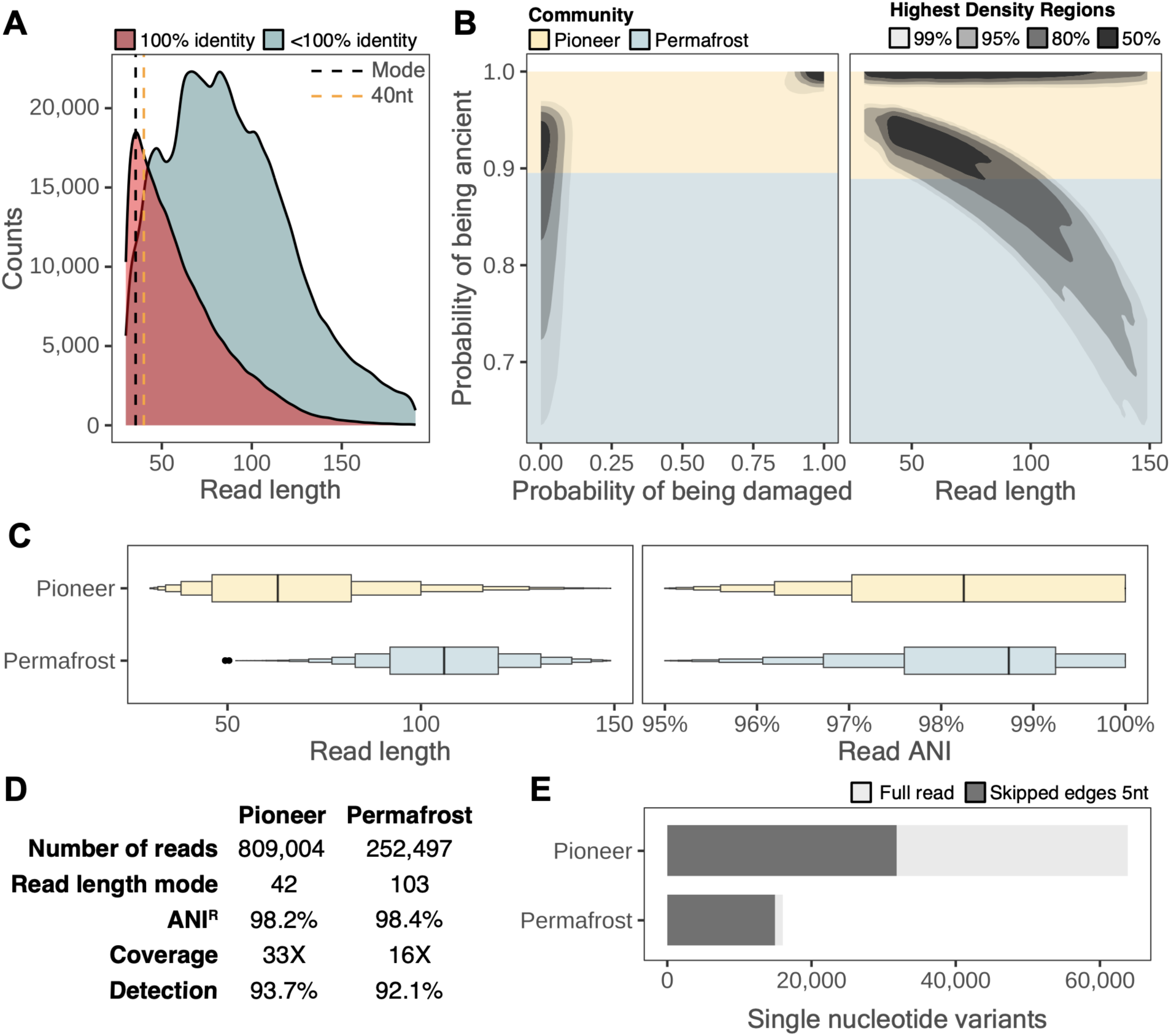
The Kap København “time-travelling” microbe. A, Read length distribution of the reads mapping to the reference 3300025461_7 from the genus g Bog-38 in sample 119_50. The orange dashed line depicts the threshold we used to exclude the sequences mapping to the reference at 100%. B, The Highest Density Regions plot shows the relationship between the probability of being ancient and the probability of being damaged and the read length. We used a probability of being ancient of 0.89 as the threshold to split the reads between pioneer and permafrost communities. C, Letter-value plots depicting the read length and read Average Nucleotide Identity to the reference 3300025461_7 for both fractions. D, Table summarising the statistics of the mapping of each fraction to 3300025461_7. E, Number of single nucleotide variants for each fraction as inferred by anvi’o, including (light grey) and excluding 5nt on the read’s edges (dark grey).

We also investigated the association between the probability of a given read to be ancient and its length, where a gradual decrease in probability as a function of increasing read lengths is expected. To establish the ANI threshold for distinguishing read fractions associated with the pioneer or permafrost communities, we selected the lowest limit within the 50% highest density region, considering both PD and read length. We added the excluded reads due to their 100% identity to the reference to the pioneer read fraction; it is a common feature in ancient metagenomic studies that many short reads are 100% identical to the reference. The reads assigned to pioneer communities had a significantly smaller read length (mode: 42 nt) than those assigned to permafrost communities (mode: 103 nt). Despite these differences, the ANI^R^ values remained remarkably consistent (Fig. 7C). In both cases, we recovered more than 92% of the modern reference genome, with coverage of 33X and 16X for the pioneer and permafrost fractions, respectively. However, differences in coverage are unlikely quantitative since our extraction protocols prioritise the recruitment of short DNA fragments, which reduces DNA recovery from living organisms.

Lastly, we implemented a new profiling method in anvi’o^73^ to exclude the edges of short reads while calculating single-nucleotide variants (SNVs) for more accurate characterisation of genuine genetic variants in ancient DNA by minimising the impact of post-mortem changes. Using this approach, we re-profiled each fraction by excluding five nucleotides from the read edges (Fig. 7D) and found 31,802 SNVs in the pioneer community, two times more than in the permafrost (14,933 SNVs). The higher extent of genetic variation we observed in the pioneer community reflects the result of the deposition over ∼20,000 years, resulting in a blend of local and upstream organisms, while the reduction of genetic variants in the permafrost fraction demonstrates how the community is shaped by selection for survival and growth under relatively stable permafrost conditions.

## Discussion

Our study provides unprecedented insights into the structure and function of the bacterial, archaea, and viral communities in the Kap København Formation during a 20,000-year period two million years ago. By distinguishing pioneer microbial communities from those that later thrived in the permafrost, we reinforce evidence of an ice-free North Greenland with a warmer, wetland-like ecosystem during the Pliocene/Pleistocene transition. Additionally, our reconstruction of carbon processing pathways suggests that high-latitude ecosystems may have contributed to moderate methane emissions driven by the northward expansion of wetlands. This aligns with early Pleistocene ice core data, which indicate that atmospheric methane concentrations and their glacial and interglacial fluctuations were not significantly higher than those observed after the mid-Pleistocene transition. However, while Arctic wetlands may have served as an additional methane source, other processes, such as methane consumption by high-affinity methanotrophs, could have reduced emissions from these high-latitude ecosystems^66^, counterbalancing these effects.

A striking aspect of our findings is the potential re-emergence of dormant methanogens preserved in permafrost over geological timescales. These “time-travelling” microbes^2,16^, capable of reviving when conditions become favourable^28,71,74,75^, provide a compelling perspective on microbial evolution and resilience. Members of the recovered genus *Methanoflorens* appear to play a central role in methane cycling, potentially reinforcing climate feedback loops in thawing permafrost^32^. Remarkably, despite being buried for at least two million years, these methanogens show >98% nucleotide sequence identity to modern relatives that have persisted across vast temporal and spatial scales, including a 3,000 km geographical separation from their closest known modern reference^71^. This observation supports the idea that not all organisms respond to environmental change in the same way. While some may remain dormant for millennia until favourable conditions return, others evolve or are replaced by new lineages in response to environmental change^2,16^.

The recovery of pioneer microbial and viral communities, alongside those of plants and animals prior to permafrost formation, provides a more comprehensive reconstruction of the Arctic ecosystem two million years ago, when temperatures were 10C warmer. Intriguingly, while the plant and animal eDNA^1^ reveal communities that vastly differ from what we can observe in the Arctic nowadays, the pioneer microbial community shows striking similarities to those of today’s thawing Arctic. This suggests that the initial response to rising temperatures is a shift in the microbial community composition, serving as early indicators of climate-driven ecosystem transformations. It also raises the possibility that initial microbial changes similar to those seen at the Pliocene/Pleistocene transition shape the Arctic landscape for an eventual change of the plant and animal communities into something similar to what is seen at the Kap København Formation two million years ago.

### Material and methods

### Metagenomic data

We retrieved the shotgun metagenomic data from Kjær et al. (ENA project accession PRJEB55522) and pre-processed it according to the same protocol described in Kjær et al. We analysed 41 samples (53 metagenomic libraries) distributed across three stratigraphic units, B1, B2 and B3, spanning five different localities (50, 69, 75 and 119). We created a short version of the sample names used by Kjær et al. (Supplementary Table S1)

### Post-depositional uplift history of Kap København Formation

We estimated a post-depositional uplift history to determine when the shallow marine Kap København Formation emerged above sea level and became permafrozen. We adopted a steady uplift rate assumption for the broader region, which resulted from the flexural isostatic response of the solid Earth to the erosional unloading caused by the fjord incision. According to our model, the upper undisturbed sediments of the Kap København Formation were deposited at approximately 15 meters of water depth around 1.93 million years ago and subsequently uplifted to their present elevation of about 165 meters above sea level, with the different sample sites being exposed subaerially between 0.8-1.2 million years ago.

### Taxonomic database generation

We used a simplified version of the taxdb-integration workflow (https://github.com/chassenr/taxdb-integration) to compile a unified taxonomy with ten levels that combined NCBI and GTDB taxonomies for our genomic references. We sourced data from various genomic datasets, including NCBI, PhyloNorway plastid sequences, complete GTDB v207, and high-quality IMG/VR v4 genomic data. We also incorporated metagenome-assembled genomes from TARA Oceans, Genomes from Earth’s Microbiomes, the glacier microbiome catalogue, and those recovered from a permafrost study.

To ensure consistency with GTDB v207, we re-annotated the bacterial and archaeal metagenome-assembled genomes using gtdb-tk v2. Subsequently, we used derep-genomes (https://github.com/aMG-tk/derep-genomes) to dereplicate the assemblies at the species or vOTU level for viruses. We then employed the viral, archaeal, bacterial, and organelle dereplicated data to construct a Bowtie2 database. Before the database construction, we concatenated the contigs of each assembly with stretches of 50Ns.

### Functional database generation

To generate a non-redundant version of the KEGG GENES database (downloaded in February 2022), we clustered the amino acid sequences of each KEGG orthologous entry using MMseqs2 (version b0b8e85f3b8437c10a666e3ea35c78c0ad0d7ec2). We used the following parameters for clustering: *-c 0.8, --min-seq-id 0.9, --cov-mode 0, and --cluster-mode 2*. To ensure reproducibility, we developed a Snakemake workflow, available at https://github.com/aMG-tk/kegg-db-setup.

### Best search parameters estimation

To determine the optimal parameters for different searches, we used a workflow (https://github.com/aMG-tk/aMGSIM-smk) that combined our workflow and *aMGSIM* (https://github.com/aMG-tk/aMGSIM) tools to generate synthetic metagenomes. We used ten samples from the Kap København formation (Supplementary Information, section 4; Supplementary Table S18-S22) to model the community composition (bacteria, archaea, and viruses), the fragment length distribution, and the damage patterns for each reference. Each synthetic metagenome contained a maximum of 1000 damaged references (determined using Bayesian damage estimates), 500 non-damaged references, and 10 million reads. We then annotated these synthetic metagenomes using the taxonomy module in our workflow, exploring different bowtie2 parameters (-N, -k [100, 250, 500, 750, 1000]), read ANI filtering thresholds (92%, 93%, 94%, 95%, 96%), and breadth filtering (Supplementary Table S18-S22). We evaluated the sensitivity and specificity of the taxonomic profiling, the abundance and damage estimations using precision, recall, F1, and F05 for classification problems, and Spearman correlation and median absolute error for quantitative comparisons.

### Taxonomic profiling

To recover the taxonomic profiles from ancient metagenomic data, we employed the taxonomic module of our workflow, an ancient metagenomics analysis workflow. We converted the reads into super-reads using Tadpole, a kmer-based assembler included in the BBTools^76^ software suite, with strict parameters. The resulting super-reads were then dereplicated using VSEARCH^77^ *–fastx-uniques*. We used the dereplicated reads for functional and taxonomic profiling. Then, we mapped the super-read dereplicated reference sequences to the original quality-controlled ancient reads, and using Bowtie2, our workflow mapped the reads against the database we generated for this study. We removed mapping duplicates using the MarkDuplicates program from Picard^78^, and filtered the BAM files with the *filterBAM* program (https://github.com/aMG-tk/bam-filter) using the following parameters: *-N -g auto -e auto -n 100 -b 0.75 -c 0.4 -A 94 -a 90 --read-length-freqs--read-hits-count --include-low-detection --min-breadth 0.01*. Finally, we estimated the post-mortem damage using Bayesian estimates in metaDMG in local and LCA modes. For the specific versions of the program used, check the files at https://github.com/GeoGenetics/2025-kapk-microbial

### Abundance table generation

We used *filterBAM* to estimate the taxonomic abundances based on the 80% truncated average depth (TAD80). Briefly, we estimated the number of reads in the TAD80 region using the formula N = (G * C) / L, where G stands for the length of the region where we estimated the TAD80, C for the TAD80 coverage value and L for the average read length mapped to the reference. We then estimated the taxonomic abundance by normalising the number of reads by the length of the region where we estimated the TAD80 and scaled by one million.

As we used the *--include-low-detection* parameter in *filterBAM*, there will be cases when the TAD80 estimated taxonomic abundance is 0. In this case, we used the taxonomic abundances estimated by the number of reads mapped normalised by the reference length and scaled by 1 million.

### Damage threshold estimation

To determine the intervals of post-mortem damage and the minimum number of reads required for reliable damage estimates of non-eukaryotic references, we used metaDMG to estimate damage in local mode and with a significance threshold of at least 2, focusing on the estimates for chloroplasts and mitochondria references (Extended Data Figure 4A). We used letter-value plots^79^ to analyse the lower tails of the damage estimate distribution for all samples and references. Subsequently, we considered only those references classified as damaged in all downstream analyses.

### Contamination identification and removal

We followed the same processing procedure for the 13 negative extraction and library controls as for the Kap København Formation samples (Supplementary Table S2). For each reference found in a control, we assessed the number of mapped reads and their damage patterns and excluded them from further analysis using an approach similar to that of Kjær et al.^1^.

### Microbial source tracking analyses

To create the source dataset for the source tracking analyses, we used *getBiomes* (https://github.com/aMG-tk/get-biomes) to retrieve the biome information and associated raw sequence data from MGnify and ENA. We ran getBiomes with the parameters *--ena-filter {library_layout: PAIRED, library_strategy: WGS, library_source: METAGENOMIC, library_selection: RANDOM, read_count: 10000000}’ --combine --exclude-terms human,16S*. Raw sequences were then processed using the QC module in our workflow with modern metagenomics settings. This involved using fastp to filter low-quality reads and selecting the forward pair for any read pairs that did not merge. The resulting metagenomes were then taxonomically annotated using our workflow with modern metagenomics settings, which turned off the read-extension step and set the read ANI to 95%.

We used the same database and procedure to annotate, filter, and estimate the taxonomic abundances of the sinks. We combined the taxonomic annotations of the sources and sinks. We filtered the combined dataset by excluding samples with fewer than 10,000 counts and species observed fewer than three times in at least 20% of the samples, with a coefficient of variation of less than three and a mean proportion across all samples of less than 1e-5. This filtering step was undertaken to ensure that only consistently present and abundant species in the samples were included in the final dataset. The resulting tables were exported as BIOM objects that were used as input for meta-Sourcetracker using the parameters *--sink_rarefaction_depth 0 --source_rarefaction_depth 0 --per_sink_feature_assignments --restarts 100 --draws_per_restart 5 --diagnostics*.

We used decOM with default parameters as a complement to meta-Sourcetracker. To create a custom k-mer matrix, we employed the kmtricks pipeline, using the same sources as meta-Sourcetracker. The parameters we used for kmtricks were *--kmer-size 29 --mode kmer:pa:bin --nb-partitions 20000 --restrict-to-list 1000 and --recurrence-min 3*.

### Extraction of reads from damaged references

We used *dReads* (https://github.com/aMG-tk/dmg-reads) to separate the reads into different domains (Eukarya, Bacteria, Archaea, and Viruses) and classify them based on their damage status. For the read extraction, we used the parameters *-f ’{ damage: 0.11, significance: 2}’ --fb-filter ’{breadth: 0.01, n_reads: 100}’ --rank ’{domain:[d Bacteria, d Archaea, d Viruses, d Eukaryota]}’*. The number of reads and damage values were estimated based on our analysis of Eukaryotic damage estimates.

### Biome-associated metagenomic read-recruitment analyses

To investigate the distribution of reads found in damaged references across multiple biomes, we constructed two additional bowtie2 databases. The first database consisted of MAGs from the Genomes from Earth’s Microbiomes catalogue^27^ and the glacier microbiome catalogue^23^. For the second database, we aimed to obtain a comprehensive overview of a permafrost thaw gradient (palsa, bog, fern) by utilising MAGs recovered from Woodcroft et al.^28^. We employed the same mapping, filtering (expected breadth ratio >= 0.25), and damage estimation strategy used in our previous analyses.

### Lipid biomarker and isotope analyses

Ancient eDNA results are supported by the analysis of microbial lipid biomarkers and their carbon isotopic compositions for a subset of ten samples. For each of these, about 10 g of sediment was extracted following a modified Bligh & Dyer procedure: Samples were ultrasonicated and extracted into a solvent mixture of methanol, dichloromethane (DCM), and aqueous buffer (2:1:0.8). A phosphate buffer (8.7 g L^-1^ KH_2_PO_4_, pH 7.4) and a trichloroacetic acid buffer (50 g L^-1^, pH 2) were each used twice. The sonicated sediment was centrifuged, the supernatants were combined in a separatory funnel and DCM and milliQ water were added for phase separation. After transferring the organic phase, the aqueous phase was extracted three more times with DCM. The obtained total lipid extract (TLE) was dried under a gentle stream of nitrogen and stored at -20°C.

Bacteriohopanepolyols and archaeal lipids were directly quantified in the TLE by high-performance liquid chromatography-mass spectrometry (HPLC-MS) following the protocol developed by Hopmans et al.^42^. A maXis quadrupole time-of-flight MS (Bruker Daltonik) equipped with an electrospray ionisation source and coupled to a Dionex Ultimate 3000RS UHPLC was employed. MS detection was carried out in positive ionisation mode while scanning a *m*/*z* range from 150 to 2000. MS^2^ scans were obtained in data-dependent mode and with the addition of an inclusion list featuring the exact masses of known BHPs. Compounds were identified according to exact precursor mass to charge (*m*/*z*), retention time and fragmentation patterns.

Additionally, for one selected sample (69_34 in unit B2), the carbon isotopic compositions of archaeol and diplopterol were obtained. From 10% of the dried TLE, *n*-hexane-soluble maltenes were recovered and subsequently separated into fractions of increasing polarity using a SUPELCO LC-NH_2_ glass cartridge. Compounds were analysed as trimethylsilyl ethers after reaction with *N,O*-bis(trimethylsilyl)trifluoroacetamide (BSTFA) in pyridine at 70°C for half an hour and identified on a Thermo Scientific ISQ gas chromatography-MS (GC-MS) system using a 30 m Restek-5ms column (0.25 mm internal diameter, 0.25 µm film thickness). The initial GC oven temperature was held at 100 °C for 1 min, increased to 150 °C at a rate of 10 °C min^−1^, then to 320°C at a rate of 4 °C min^−1^ and held for 41.5 min. Compound specific isotope analysis was performed on a Trace GC Ultra coupled via a GC Isolink interface to a Delta V plus isotope ratio MS (Thermo Fischer Scientific) using the same temperature program as for GC-MS analysis. Stable carbon isotope values are reported in the delta notation as δ^13^C values relative to Vienna Pee Dee Belemnite with a precision of better than 1‰ based on repetitive measurements of an in-house *n*-alkane standard. These values were corrected by mass balancing for the contribution of carbon atoms added during derivatisation.

### Functional profiling

We used the functional profiling module of our workflow to annotate the dereplicated super-reads with MMseqs2, using fine-tuned parameters for ancient DNA amino acid searches (*--comp-bias-corr 0 --mask 0 -e 1e-5 --exact-kmer-matching 1 --sub-mat PAM30.out -s 3 -k 6 --spaced-kmer-mode 1 --spaced-kmer-pattern 11011101 --min-length 15 --format-mode 2 -c 0.8 --cov-mode 2 --min-seq-id 0.6*). To perform the annotation, we used the non-redundant KEGG GENES database and dbCAN2 v11 gene sequences (CAZyDB.08062022). We applied *xFilter*, which implements a modified version of FAMLI to identify the KEGG/dbCAN2 genes most likely present in the samples with the parameters *-n 25 -b 20 -e 1e-5 --breadth-expected-ratio 0 -f depth_evenness --depth-evenness 1*. For KEGG-related results, xFilter produced output compatible with the enzymes-txt mode of anvi-estimate-metabolism in anvi’o.

We expanded the standard KEGG modules collection in anvi’o using anvi-setup-user-modules to include custom modules (Supplementary Table S12; defined as Woodcroft2018 and Borrel2023 in the module class) designed to explore carbon metabolism in permafrost. We ran anvi-estimate-metabolism with the parameters *--add-coverage --output-modes modules,module_paths,module_steps,hits --include-kos-not-in-kofam --user-modules*, and we parsed the output to identify pathways with 100% completeness.

Furthermore, we extracted hits to CAZy families related to cellulose degradation (GH5, GH9, 3.2.1.4, GH51, GH6, GH7, GH48, 3.2.1.91), xylan degradation (GH5, GH8, GH10, GH11, GH43, 3.2.1.8, GH3, GH30, GH39, GH52, GH54, GH116, GH120, 3.2.1.37, GH67, GH115, 3.2.1.139, CE1, CE2, CE3, CE4, CE5, CE6, CE7, CE12, 3.1.1.72), and xylose degradation (3.2.1.37, GH3, GH30, GH39, GH10, GH43).

### Reference metabolism estimation

We used the program *anvi-estimate-metabolism* from anvi’o with default parameters to estimate the metabolisms of the archaeal and bacterial references that recruited reads in our study. Note that we used a version of KEGG downloaded in January 2023 (for reproducibility, the hash of the KEGG snapshot available via *anvi-setup-kegg-kofams* is d20a0dcd2128)

### Virome characterisation

To increase the sensitivity of the viral taxonomic classification, we envisioned an approach to identify which viral reference is most likely present in a sample. We created a protein database containing 15,948,451 sequences from the viral sequences from IMG/VR v4 present in the database we used for the nucleotide-based taxonomic profiling, and we enriched it with 16,441 proteins from a curated collection of mobile genetic elements (MGE) associated with methanogenic archaea. We used MMseqs2 with the fine-tuned parameters for ancient DNA amino acid searches (*--comp-bias-corr 0 --mask 0 -e 1e-5 --exact-kmer-matching 1 --sub-mat PAM30.out -s 3 -k 6 --spaced-kmer-mode 1 --spaced-kmer-pattern 11011101 --min-length 15 --format-mode 2 -c 0.8 --cov-mode 2 --min-seq-id 0.6*) and then filtered the results with *xFilter* to identify the most likely proteins present in the samples with the parameters *-n 25 -b 20 -e 1e-5 --breadth-expected-ratio 0 -f depth_evenness --depth-evenness 1*. Then, we estimated the proteome completion ratio for each reference and selected all the ones with a value over 10%. We functionally annotated the selected proteins using HHblits using two iterations in combination with PFAM v35 and PHROG v4. We followed a similar approach to that of Vanni et al.^80^ to remove overlapping domains and select those hits with a probability over 90% and a target or query coverage larger than 40%. We manually curated the results and classified them into six functional categories: “Unknown”, “Virus structural protein”, “DNA/RNA transactions/processing”, “Lysis”, “TA/RM systems” and “AMG and other”.

### Estimation of probabilities of being ancient and being damaged

First, we extracted and aligned reads from damaged references against the taxonomic profiling database. Unlike previous alignments, we did not use the -k option in bowtie2 and reported the best alignment. The filtering and damage estimation steps were conducted as described earlier. We then used the program *getRPercId* (https://github.com/aMG-tk/get-read-percid) to subset the BAM files to the reference 3300025461_7 and calculate for each the read length and average nucleotide identity (ANI). After exploring the read length and ANI, we subset the BAM files to reference 3300025461_7 and calculate the read length and average nucleotide identity (ANI) for each read. Then, for a more formal characterisation of the deamination patterns observed in each ancient DNA strand for each fraction of data, we estimated the Briggs parameters, namely ***δ***d (deamination rates within the double-stranded regions), ***δ***s (deamination rates within the single-stranded regions), ***v*** (nick occurring rate in the sample), and ***λ*** (overhang length distribution) using ngsBriggs. The method is based on the counts of nucleotide changes (including but not limited to C to T and G to A) at the first and last 15 cyclic positions of the strands and the position-specific error rates. The tool conducts a multinomial regression to estimate the four parameters (***δ***d, ***δ***s, ***v*** and ***λ***) that describe the deamination pattern of the ancient DNA samples. Under the assumption that the length distributions of the ancient pioneer strands and modern contaminated strands are distinguishable, the tool will then go through each strand within the length range [30,150) and estimate the best-fit ancient and modern length distributions if the prior knowledge of the modern contamination rate of the sample is provided. The previous four estimates can help to calculate the deaminated probability conditioned on the focal strand being ancient (AN) and the posterior probability of being ancient for each focal strand, given its information on nucleotide changes and fragment length (PD). We used the following parameters for the inference *-isrecal 1 -model b -eps 0.05 and -eps 0.1*. After identifying the pioneer and permafrost fractions, we estimated the Briggs parameters. Lastly, we used a new profile mode in anvi’o, specially designed for ancient DNA, to infer the SNVs in each fraction where we exclude a certain number of positions from the read edges using the option *--skip-edges 5*.

## Acknowledgements

The authors thankfully acknowledge the computer resources and the technical support provided by the BMBF-funded de.NBI Cloud within the German Network for Bioinformatics Infrastructure (de.NBI) (031A537B, 031A533A, 031A538A, 031A533B, 031A535A, 031A537C, 031A534A, 031A532B). R.L-P. was funded by a Ramón y Cajal grant (RyC2021-031775-I) from the Spanish Ministerio de Ciencia e Innovación (MCIN/AEI/10.13039/501100011033) and the European Union («NextGenerationEU»/PRTR). EW thanks the Lundbeck Foundation, the Carlsberg Foundation, the Danish National Research Foundation, the Novo Nordisk Foundation, and the Wellcome Trust for financial support and the St. John’s College, Cambridge for providing an environment for scientific discussion and thought. IV acknowledges support from the National Science Foundation Graduate Research Fellowship under Grant No. 1746045. GB was supported by the French National Agency for Research Grants (Methevol, ANR-19-CE02-0005-01). M.S. acknowledges support from the National Research Foundation of Korea (grants 2021R1C1C102065), the Samsung DS Research Fund, and the Creative-Pioneering Researchers Program through Seoul National University. VKP was supported by a research grant (15467) from the Danish foundation VILLUM FONDEN. YW acknowledges support from the NSFC BSCTPES Project (No. 41988101), the CAS Youth Interdisciplinary Team Fund, National Key Scientific and Technological Infrastructure project “Earth System Numerical Simulation Facility” (EarthLab, 2023-EL-ZD-000111), National Supercomputer Center in Wuxi, and SMU’s Center for Research Computing. Additional support was provided by Germany’s Excellence Strategy (EXC-2077), project 390741603 “The Ocean Floor – Earth’s Uncharted Interface”. The authors thank Rayan Chikhi and Camila Duitama González for their insightful discussions on effectively utilising decOM and Nadine T. Smit for guidance in the analysis of bacteriohopanepolyols.

## Data availability

All sequencing data used in this study are accessible at the European Nucleotide Archive under accession PRJEB55522. The re-extracted sequence data, taxonomic database, and other supplementary data can be found at ERDA with the following DOIs: https://doi.org/n59w, DOI: https://doi.org/n6ks, and DOI: XXX respectively.

## Code availability

All code used in this study is available at https://github.com/GeoGenetics/2025-kapk-microbial and archived on ZENODO (DOI:XXX). The repository contains a reproducible workflow for generating the basic data, as well as the analytical pipeline that produced all tables and figures. The various tools included in the ancient metagenomic toolkit can be accessed at https://github.com/aMG-tk

## Extended Data figures

**Extended Data Fig. 1.**
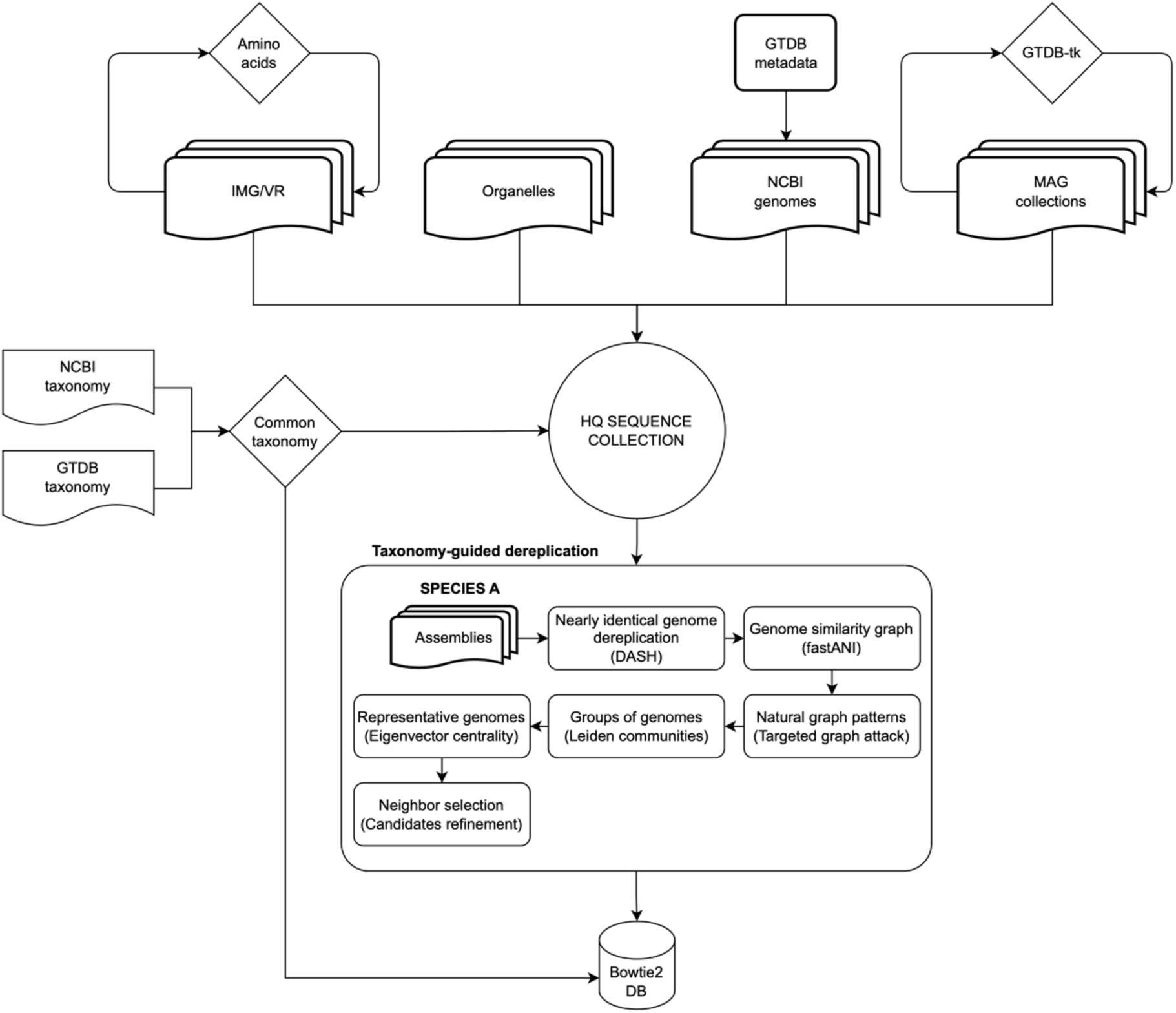
Taxonomic database creation workflow. Diagram illustrating the workflow to create the taxonomic database used for profiling the samples in this study.

**Extended Data Fig. 2.**
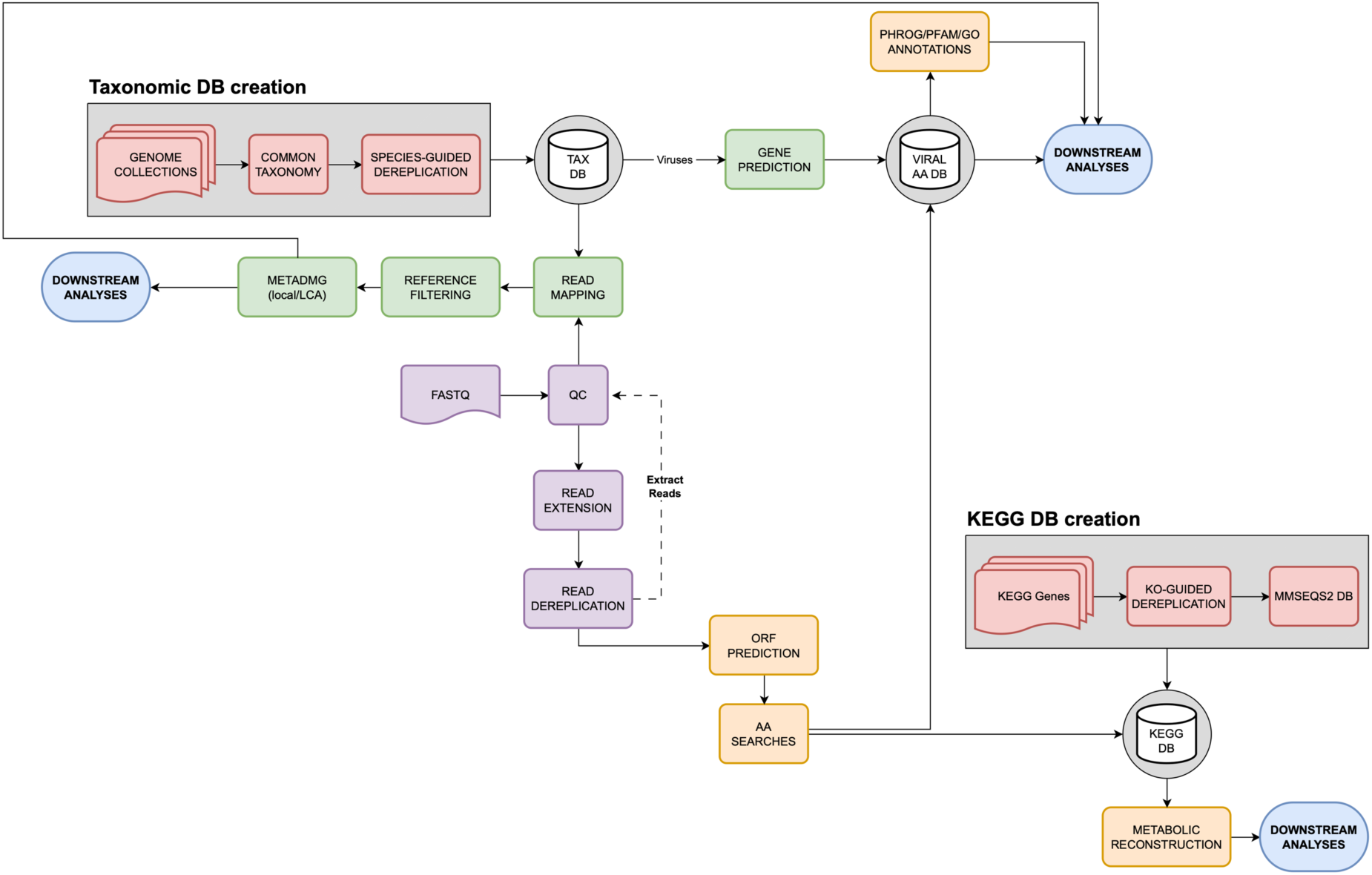
Taxonomic and functional profiling workflow. Schematic diagram illustrating the main steps implemented in the taxonomic and functional analysis workflow used to process the samples in this study. In red are the steps where databases are created; in purple, the QC post-processing steps involved in creating super-reads and dereplication; in green, the steps for taxonomic profiling; and in orange, the functional annotation steps.

**Extended Data Fig. 3.**
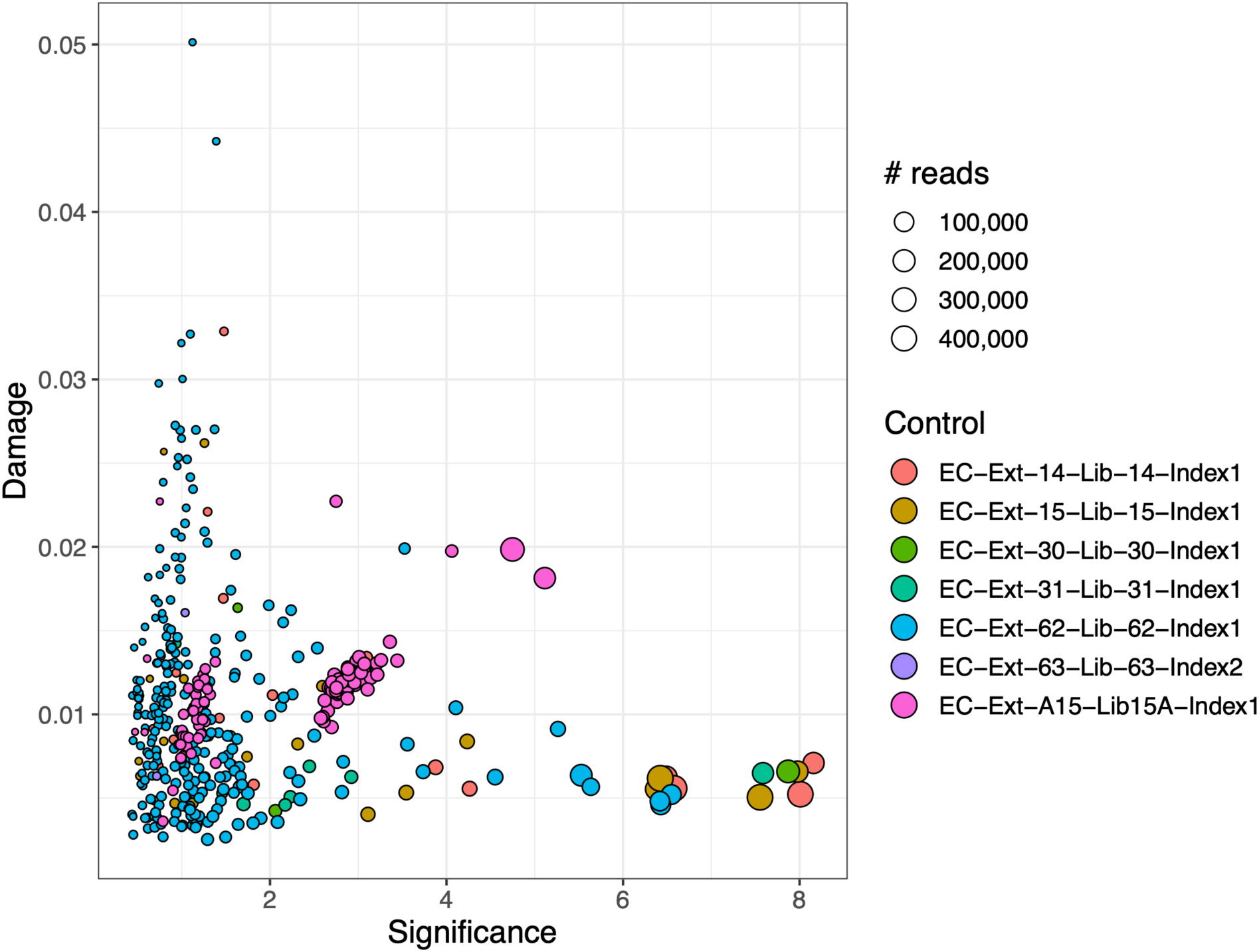
Extraction controls. Each point in the plot corresponds to a taxon identified in the control samples based on damage and its significance. We did not detect any taxa with damage. The size of the point corresponds to the number of reads mapped, while the colour indicates the control sample (Supplementary Table S2).

**Extended Data Fig. 4.**
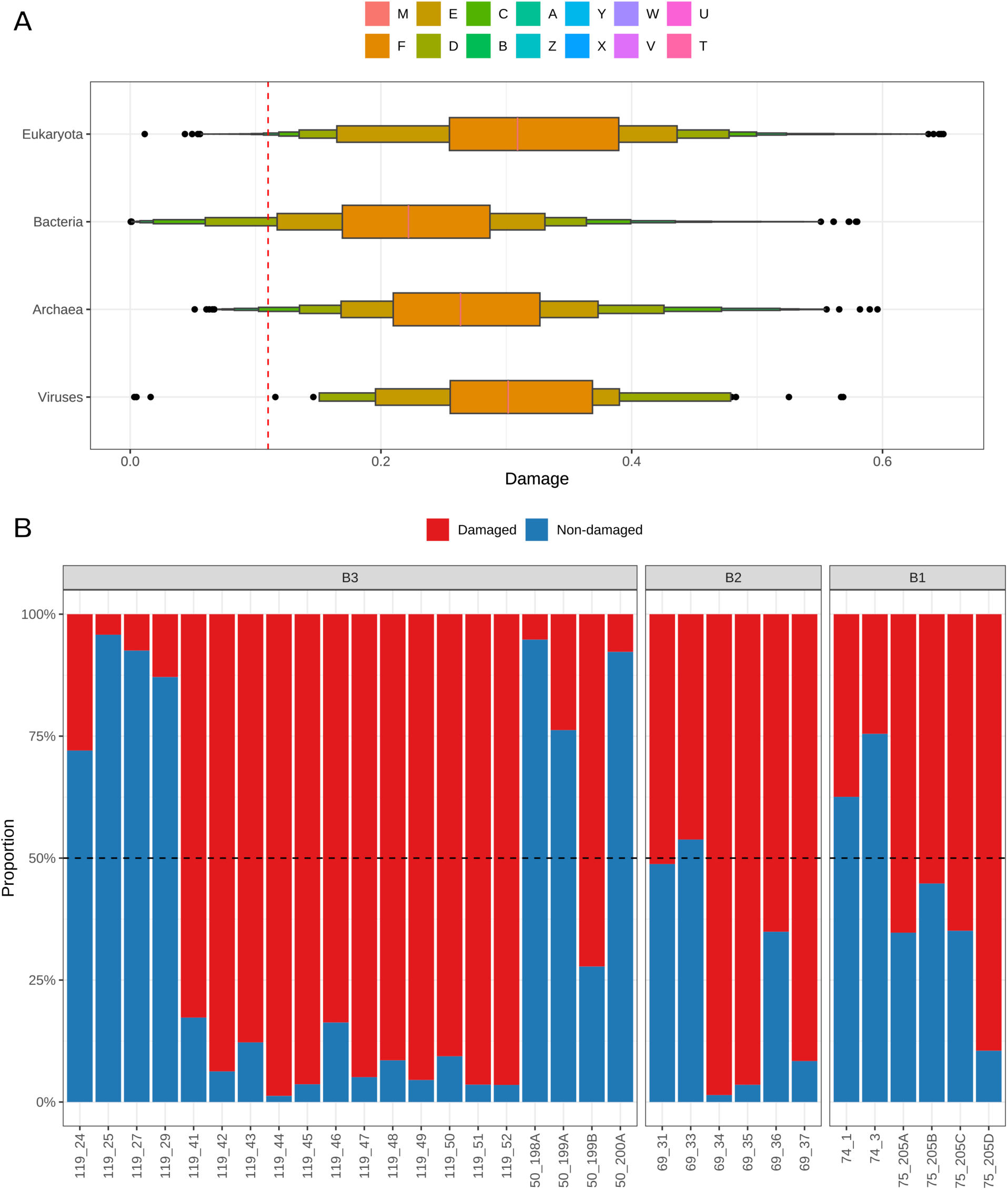
Damage thresholds and sample selection. A) Letter-value plots illustrating the damage distribution for Archaea, Bacteria, Viruses, and Eukaryotes. The red line indicates the threshold to determine whether a taxon is potentially damaged. B) We discarded those samples that had less than 50% of the estimated TAD80 taxonomic abundances.

**Extended Data Fig. 5.**
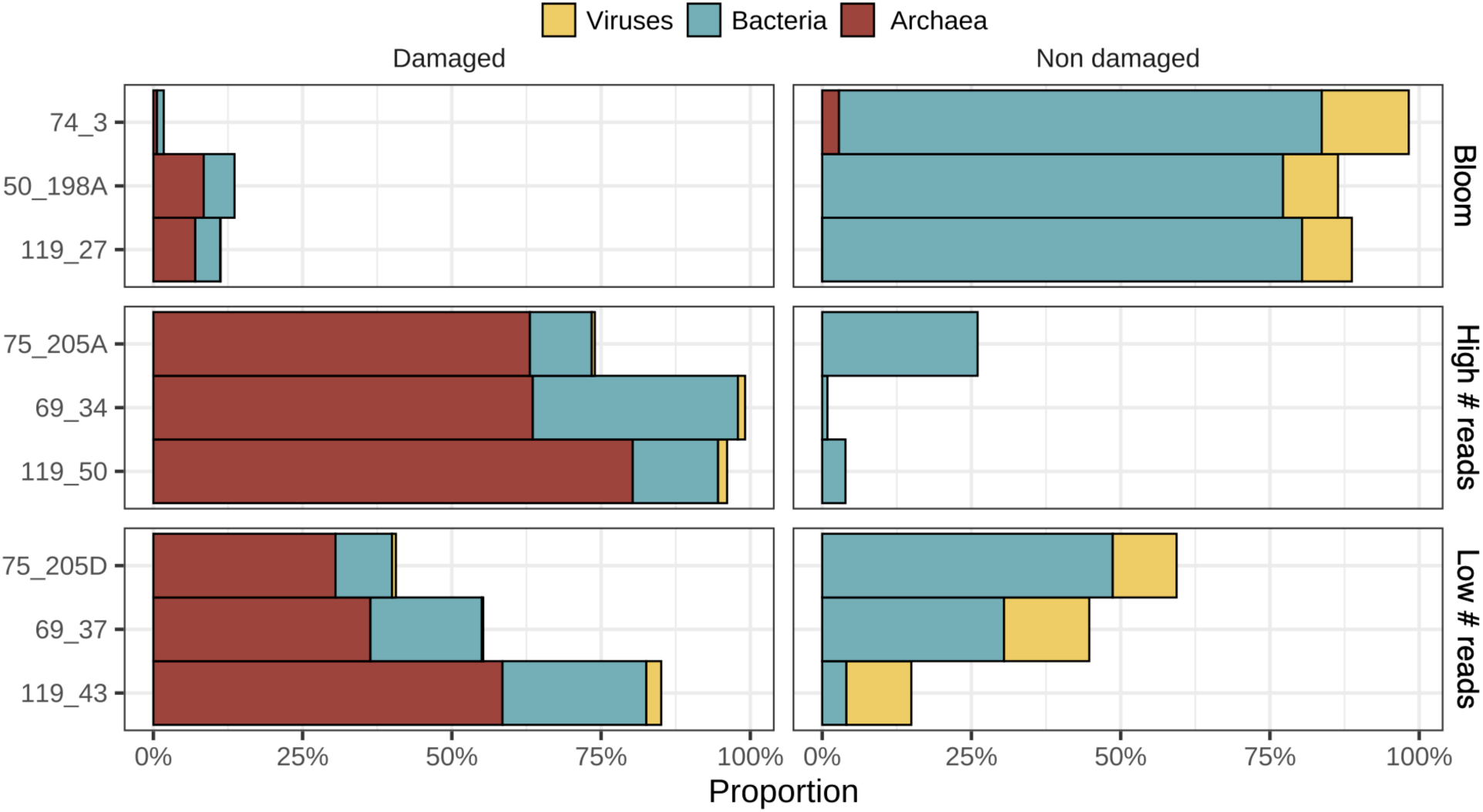
Extraction bias and bloom assessment. Proportion of different taxa after re-extracting selected samples to assess potential extraction bias and blooming of taxa. We selected three distinct sets of samples: three samples enriched in taxa that do not show damage and six samples where Archaea are abundant and produced either a high or low number of reads in the original dataset from Kjaer et al., 2022.

**Extended Data Fig. 6.**
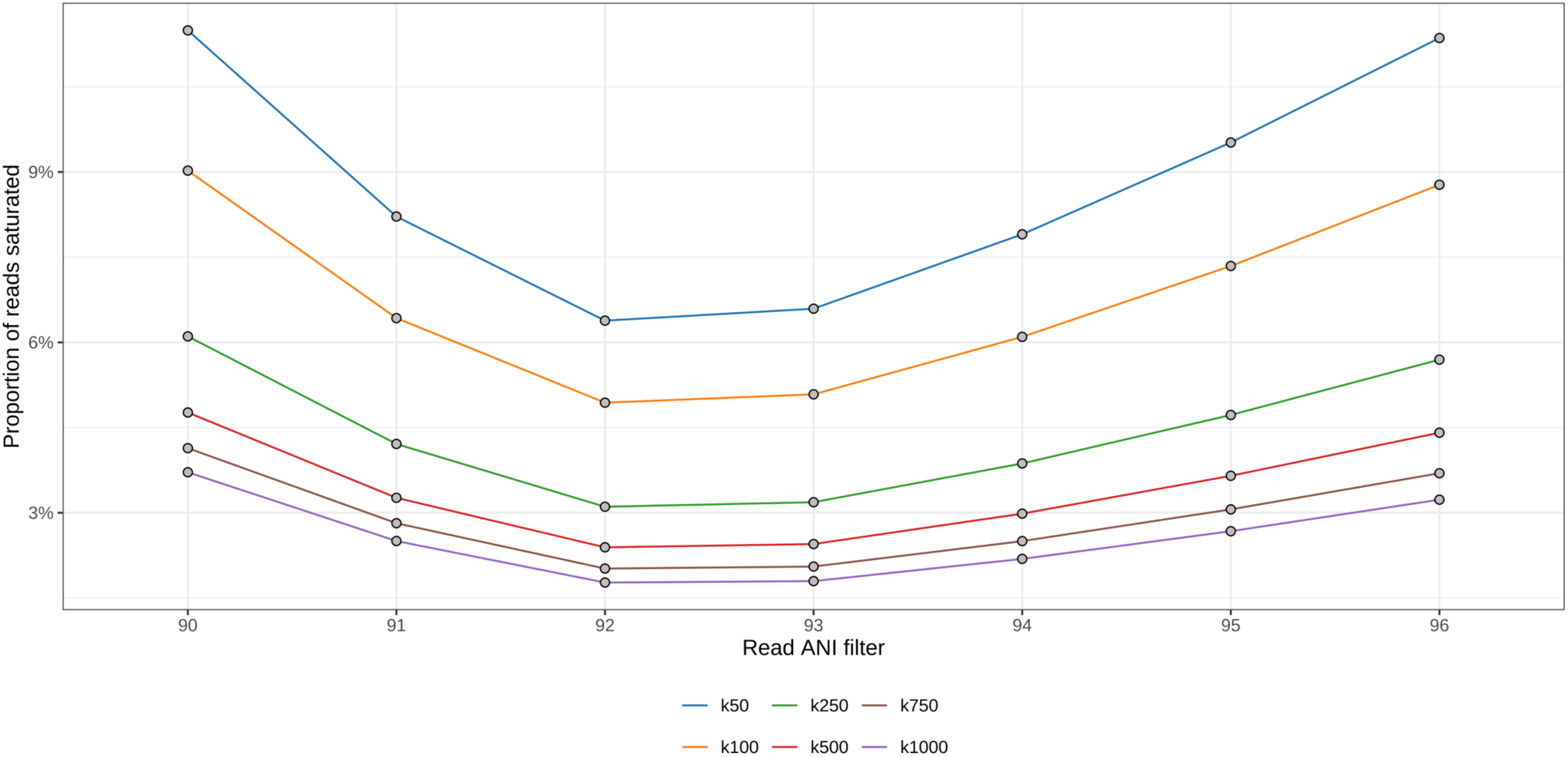
Effect of read alignment saturation. We explored various values (50, 100, 250, 500, 750, and 1000) for the Bowtie2 parameter -k, determining the number of alignments reported for each read. Our evaluation spanned multiple Average Nucleotide Identity values (90-95%). The lower the values, the better.

**Extended Data Fig. 7.**
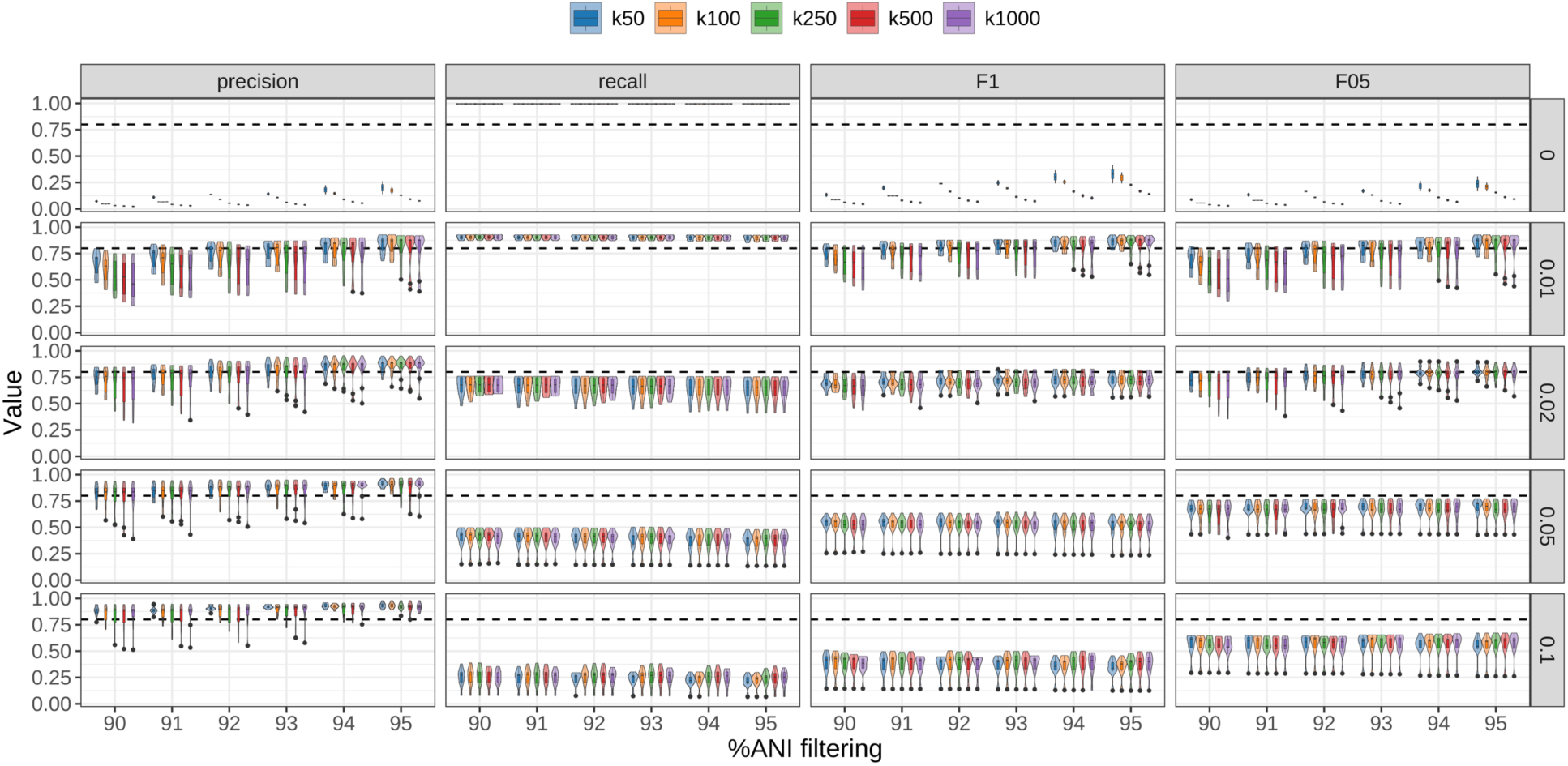
Assessment of the taxonomic classification. To determine the optimal parameter combination that accurately recovers the correct taxonomic composition of the samples in this study, we examined several values (50, 100, 250, 500, 750, and 1000) for the Bowtie2 parameter -k, alongside filtering thresholds for ANI (90-94%) and breadth of coverage (0-10%) across 10 synthetic ancient metagenomes derived from the samples used in this study. We computed Precision, Recall, F1, and F0.5 for each parameter combination. The higher the values, the better.

**Extended Data Fig. 8.**
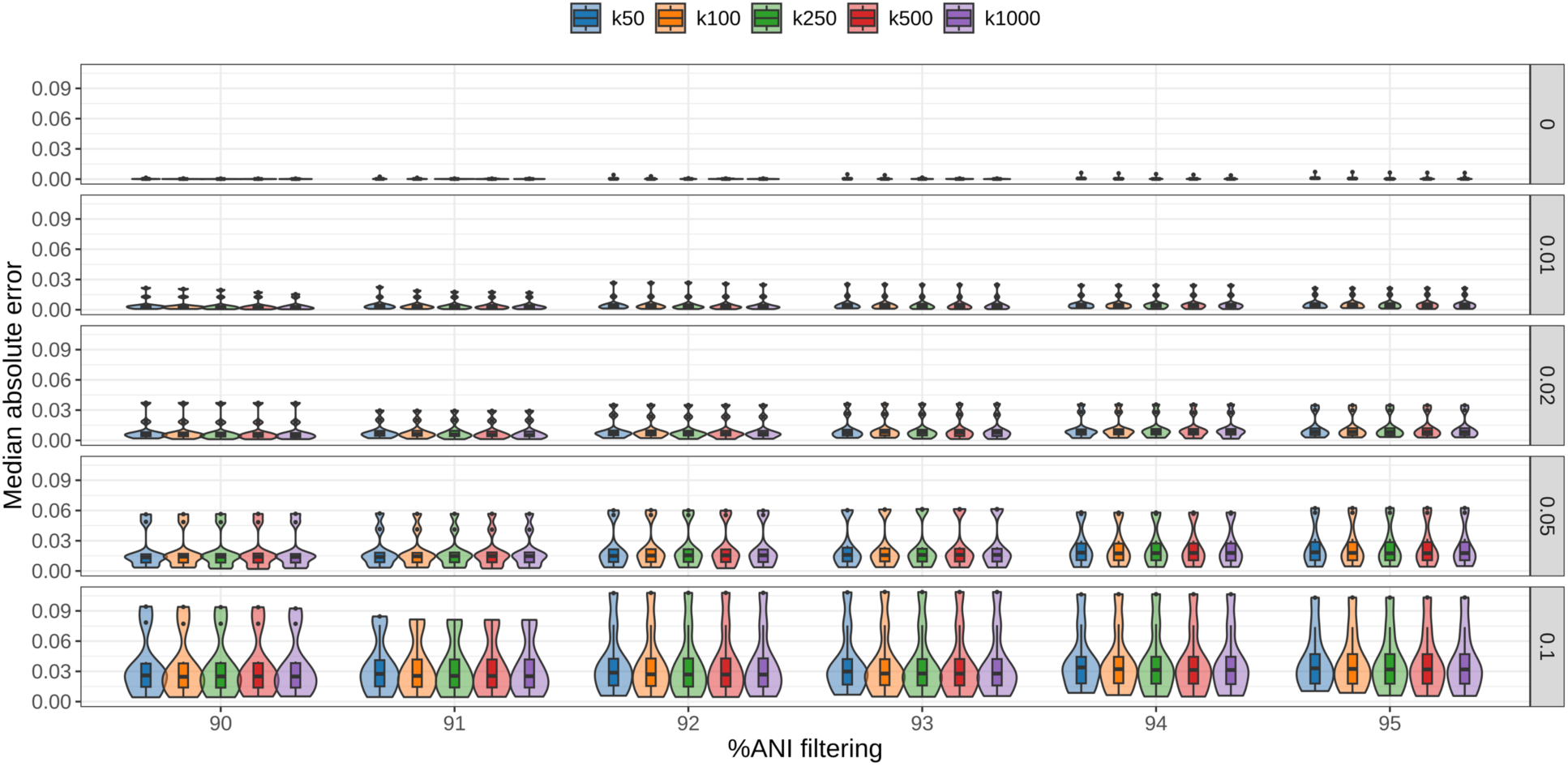
Assessment of the taxonomic abundances. To determine the optimal parameter combination that accurately recovers the correct taxonomic abundances of the samples in this study, we examined several values (50, 100, 250, 500, 750, and 1000) for the Bowtie2 parameter -k, alongside filtering thresholds for ANI (90-94%) and breadth of coverage (0-10%) across 10 synthetic ancient metagenomes derived from the samples used in this study. For each parameter combination, we computed the Median Absolute Error. The lower the values, the better.

**Extended Data Fig. 9.**
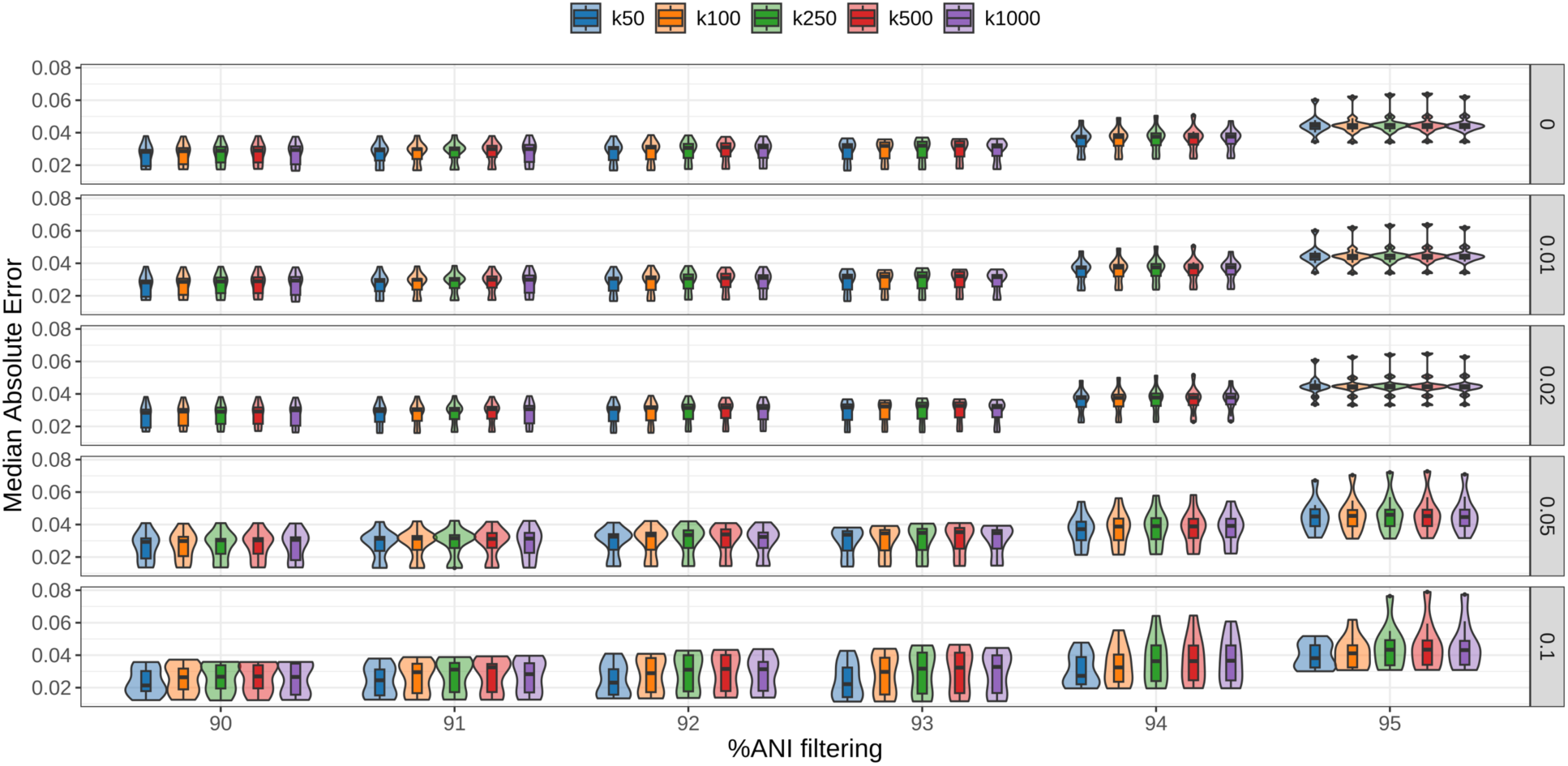
Assessment of the damage estimates. To determine the optimal parameter combination that accurately recovers the correct taxa damage estimates of the samples in this study, we examined several values (50, 100, 250, 500, 750, and 1000) for the Bowtie2 parameter -k, alongside filtering thresholds for ANI (90-94%) and breadth of coverage (0-10%) across 10 synthetic ancient metagenomes derived from the samples used in this study. For each parameter combination, we computed the Median Absolute Error. The lower the values, the better.

