## Supplementary Information for "Two-million-year-old microbial communities from the Kap København Formation in North Greenland"

### 1 Vegetation in the Kap København formation

Among the abundant tree species identified in the Kap København Formation by the ancient eDNA were poplars (*Populus*). While the natural range of Balsam poplars (*Populus balsamifera*) commonly is below the treeline, it can today be found on the northern slope of the Brooks Range, an area characterised by continuous permafrost. However, these poplar groves are confined to river channels within a limited range of a few hundred meters. These areas often exhibit a 'thaw bulb,' a depression in the permafrost table[1]. Other genera found in the samples, such as *Crataegus*, *Rhamnus*, *Thuja*, and *Kalmia*, are distributed in northern temperate regions. However, they are not adapted to boreal environments and cannot survive, even as clones, in a permafrost-affected environment. In particular, *Kalmia angustifolia* and *Thuja occidentalis* are typically found in mixed deciduous forests of the Appalachia region, extending north to the southern boreal margin in Quebec. They cannot withstand Arctic winters or grow on permafrost. Hence, these genera must have colonised northern Greenland by dispersing through the Baffin and Ellesmere islands when the region was free of continuous permafrost. This suggests that they may have migrated northeast during a relatively warmer interval.

### 2 Extraction bias and bloom assessment

To assess the potential effect of extraction bias responsible for the observed high archaeal abundances and to evaluate a possible bloom caused by sample handling, we optimised the extraction protocol for ancient environmental DNA to re-extract nine samples that fell under one of these categories:

- Bloom: Three samples where we observed a relative abundance > 50% of taxa showing no damage.
- High number of reads: Three samples with a large relative abundance of archaeal taxa and generating a large number of reads (high DNA yield).
- Low number of reads: Three samples with a large relative abundance of archaeal taxa and generating a low number of reads (low DNA yield).

We supported the existing results in all cases, as shown in Supplementary Table S5-S6 and Extended Data Fig. 5.

### Methods

DNA was extracted from 0.3 g of sediment using the combination of solutions used in two extraction soil kits, FastDNA™ Spin Kit for Soil (MP Biomedicals) and Magattract PowerSoil Pro DNA kit (Qiagen), following the protocol we designed to extract short DNA fragments down to 30 bp to the PowerBead Pro Tube (Qiagen) containing sediment, we added 978 µL Sodium Phosphate buffer (MP Biomedicals) and 122 µL MT buffer (MP Biomedicals). We homogenised the tubes on FastPrep® instrument for 20 seconds at 4.5m/s speed and incubated the samples on the rotator for 25 min at room temperature. We centrifuged the tubes at 13000 x g for one minute and transferred the lysate to a new 1.5 mL LoBind tube (Eppendorf). Next, we added 250 µL protein precipitation solution PPS (MP Biomedicals) to the lysate, mixed and centrifuged at 13000 x g for one minute. Afterwards, we added 300 µL of CD2 inhibitor removal solution (Qiagen) and mixed the tubes thoroughly. We centrifuged the tubes again at 13000 x g for one minute and transferred the lysate to a new tube.

We transferred 450 µL lysate to the tube containing 30 µL silica magnetic beads (G-Biosciences) and 1450 µL modified QSB1 binding buffer (Qiagen). The binding buffer was modified to ensure the binding of short DNA fragments. After adding isopropanol to the QSB1 buffer according to manufacturer instructions, we added 3M sodium acetate (pH 5.2) (Sigma Aldrich) to the final concentration of 0.15 M and 5M sodium chloride (Thermo Fisher) to the final concentration of 0.025M.

We incubated the tubes on a rotator at room temperature for 15 minutes. Afterwards, we placed the tubes on the magnetic rack, discarded the supernatant, and washed the beads in 500 µL MW1 buffer (Qiagen), followed by two washes in 500 µL 80% ethanol. After drying the beads,

DNA was eluted in 50 µL EB elution buffer (Qiagen). The DNA extracts were converted to double-stranded libraries (Meyer and Kircher 2010) and sequenced on the Illumina NovaSeq 6000 instrument.

### 3 Methane flux estimates during the Arctic expansion in the Pleistocene

We have shown the resemblance of the ancient Kap København microbial ecosystem to modern boreal palustrine wetlands, as well as the dominance of methanogenic archaea in it. This finding suggests that such ecosystems extended further north than nowadays. To assess the potential impact this extension could have had on global $CH_4$ budgets during early Pleistocene interglacials (and potentially in the future), we estimated the resulting additional emissions that would arise if such boreal wetlands populated the high Arctic and compared them to modern wetland emissions.

We used combined $CH_4$ fluxes for fens, bogs and marshes presented by Kuhn et al.[2] as a modern reference point for our calculations. Emissions in these wetlands range between 3.4 and 17 g $CH_4$ $m^{-2}$ $yr^{-1}$ ($25^{th}$ and $75^{th}$ quartile), with a median value of 8.3 g $CH_4$ $m^{-2}$ $yr^{-1}$. These values are in good agreement with fluxes (4.3 and 6.2 g $CH_4$ $m^{-2}$ $yr^{-1}$) obtained by dividing bottom-up and top-down estimates for wetland emissions north of >60°N[3] by a long-term maximum wetland area of 2.1 million $km^2$ in the same region[4], and also with data for the mixed Stordalen mire (7.7 g $CH_4$ $m^{-2}$ $yr^{-1}$)[5].

Considering the position of Kap København at the northernmost tip of Greenland, we assumed an extreme scenario involving a potential expansion of these wetlands to the entire Arctic region currently occupied by tundra and rocklands[6]. Glaciers were excluded from this calculation. Finally, we needed to estimate which fraction of these regions would have been occupied by wetlands. A relative land cover of 15% was obtained as a modern reference by comparing current areas of fens, bogs, and marshes to the one of boreal forests[6].

With these assumptions, emissions derived from an expansion of these wetlands into the high Arctic were calculated. After subtracting current emissions by tundra wetlands and permafrost bogs[7,8], our calculations reveal additional emissions of 5.4 Tg $CH_4$ per year (ranging from -1 to 16 Tg $CH_4$ per year for $25^{th}$ and $75^{th}$ quartile $CH_4$ emissions). Compared to top-down and bottom-up estimates for modern wetland emissions[3], this corresponds to 41 and 59% of emissions from northern high latitudes (>60°N) and 3 and 3.6% of global wetland emissions.

These calculations only represent emissions derived from ecosystem changes, not the increased methane emissions from wetlands and lakes, which might result from increasing temperatures and thawing permafrost[2].

Additional radiative forcing by $CH_4$ has been invoked as one of the features explaining the underestimation of warming and polar amplification by climate models of the Pliocene and early Pleistocene[9]. Here, we present one potential source of such enhanced emissions from the high Arctic. Our estimate of a relatively moderate strengthening of methane emissions by northward expansion of wetlands is consistent with ice core data from the early Pleistocene, which suggests that atmospheric $CH_4$ concentration and its glacial/interglacial variability might not have been larger than after the mid-Pleistocene transition (1.2 – 0.8 Myr)[10]. Moreover, the potential emergence of wetlands in the Arctic tundra is also relevant for accounting for additional sources of greenhouse gases in the future[11].

#### Methods

To obtain a reference value for $CH_4$ flux from boreal wetlands, we used the daily fluxes for bogs, fens and marshes in Kuhn et al.[2]. These were converted into annual fluxes by assuming a 6-month growing season and combined by considering the relative land cover of these three types of wetlands[6]. Given the similarity of the microbial population and temperature regime from the Stordalen Mire (Sweden)[12] to the reconstructed ecosystem of Kap København, we were interested in the Stordalen Mire data as a further reference point for annual $CH_4$ fluxes. The total hydrocarbon flux (6.4 g C $m^{-2}$ $yr^{-1}$)[13] was corrected for a 10% contribution of non-methane volatile organic carbon[5], resulting in a value of 5.8 g carbon contained in methane ($CH_4$-C) $m^{-2}$ $yr^{-1}$, or 7.7 g $CH_4$ $m^{-2}$ $yr^{-1}$.

Current emissions from the tundra and rockland regions were estimated by multiplying median emissions from wet tundra and permafrost bogs[7] by their land cover[6]

### 4 Parameter selection using synthetic ancient metagenomes

To identify the best parameters for taxonomic profiling, we developed aMGSIM (https://github.com/aMG-tk/aMGSIM), a programme that simulates ancient metagenome reads in multiple microbial synthetic communities and utilises the properties of actual ancient metagenomes to generate synthetic datasets that can serve as ground truth. aMGSIM can model the community composition (bacteria, archaea, and viruses), the fragment length distribution, and the damage patterns for each reference. Each synthetic metagenome contained a maximum of 1,000 damaged references (determined using Bayesian damage estimates), 500 non-damaged references, and 10 million reads. We utilised this information to generate ten synthetic ancient metagenomes, allowing us to explore the parameter space of Bowtie2, identify the optimal parameters for our searches, and understand how these parameters affect such ancient samples. Subsequently, we processed these synthetic ancient metagenomes through our workflow and assessed the impact of each parameter on the recovery of the ground truth. The parameters we explored are:

- different bowtie2 parameters (-N, -k [100, 250, 500, 750, 1000])
- read ANI filtering thresholds (92%, 93%, 94%, 95%, 96%)
- coverage of breadth filtering (0%, 1%, 2%, 5%, 10%)

We evaluated the sensitivity and specificity of the taxonomic profiling, the abundance and damage estimations using precision, recall, F1, and F05[14] for classification problems, and the median absolute error (MAE) for quantitative comparisons.

First, we evaluated the impact of using a different set of parameters in bowtie2 (vsens, sens-N1, sens-N1-L20, vsens-N1), which could affect the sensitivity and running time of our mappings in three samples with varying amounts of reads (37M, 132M and 204M). We selected sens-N1 (predefined Bowtie2 sensitive parameters and allowing one mismatch in the seed) as the optimal configuration, as the fold-increase of reads mapped compared to the most sensitive settings (sens-N1-L20, vsens-N1) is marginal; however, the running time increased exponentially, being in the most sensitive setting, 40-60 times slower (Supplementary Table S18).

We then evaluated the effect of using different values of -k (the number of alignments returned for each read) and the average ANI values used for filtering to investigate the saturation of each read's mappings to the various references. This allowed us to assess the number of reads that do not map to new references because of the -k constraints. We identified -k 1000 as the optimal parameter, where only 2-4% of the reads are saturated, and 93% and 94% as potential ANI values, with 2.1% and 2.5% of reads saturated, respectively (Supplementary Table S19; Extended Data Fig. 6).

Next, we evaluated how the different parameters -k, ANI, and breadth filtering influence the recovery of the ground truth of the synthetic ancient metagenomes. We observed a decrease in the F05 values as we reduced the ANI values and increased the breadth of the coverage threshold (Supplementary Table S20; Extended Data Fig. 7). When not using any breadth of coverage filtering, all the metrics used for the evaluation were very low, indicating bad taxonomic profiling results (Supplementary Table S20; Extended Data Fig. 7). A minimum breadth of coverage of 0.01 (1% of the genome detected), combined with a -k 1000 and an ANI filtering of 94% or 95%. Besides the taxonomic composition, we also evaluated our ability to recover the TAD80 abundances by exploring the same parameters. We use the median absolute error to evaluate the impact of the different parameters (Supplementary Table S21; Extended Data Fig. 8). The most critical parameter was the breadth of coverage filtering; the higher it was, the worse our estimates became. We selected 0.01 as the final threshold.

Finally, we assessed how various parameters influenced the recovery of damage estimates. Based on the prior results, we determined that 94% ANI was the optimal value, offering a suitable balance between sensitivity and specificity while delivering an accurate reconstruction of taxonomic profiling in composition and abundances (Supplementary Table S22; Extended Data Fig. 9). The final parameters we employed included Bowtie2's sensitive predefined settings with -k 1000 and -N 1, followed by an ANI filtering set at 94% and a breadth of coverage of 0.01.
